# Path integration in large-scale space and with novel geometries: Comparing Vector Addition and Encoding-Error Models

**DOI:** 10.1101/809012

**Authors:** S. K. Harootonian, R. C. Wilson, L. Hejtmánek, E. M. Ziskin, A. D. Ekstrom

## Abstract

Path integration is thought to rely on vestibular and proprioceptive cues yet most studies in humans involve primarily visual input, providing limited insight into their contributions. We developed a paradigm involving walking in an omnidirectional treadmill in which participants were guided on two legs of a triangle and then found their back way to origin. In Experiment 1, we tested a range of different triangle types while keeping distance relatively constant to determine the influence of spatial geometry. Participants overshot the angle they needed to turn and undershot the distance they needed to walk, with no consistent effect of triangle type. In Experiment 2, we manipulated distance while keeping angle relatively constant to determine how path integration operated over both shorter and longer distances. Participants underestimated the distance they needed to walk to the origin, with error increasing as a function of the walked distance. To attempt to account for our findings, we developed computational models involving vector addition, the second of which included terms for the influence of past trials on the current one. We compared against a previously developed model of human path integration, the Encoding Error model. We found that the vector addition models captured the tendency of participants to under-encode guided legs of the triangles and an influence of past trials on current trials. Together, our findings expand our understanding of body-based contributions to human path integration, further suggesting the value of vector addition models in understanding these important components of human navigation.

**Author Summary:** How do we remember where we have been? One important mechanism for doing so is called path integration, which refers to the ability to track one’s position in space with only self-motion cues. By tracking the direction and distance we have walked, we can create a mental arrow from the current location to the origin, termed the homing vector. Previous studies have shown that the homing vector is subject to systematic distortions depending on previously experienced paths, yet what influences these patterns of errors, particularly in humans, remains uncertain. In this study, we compare two models of path integration based on participants walking two legs of a triangle without vision and then completing the third leg based on their estimate of the homing vector. We found no effect of triangle shape on systematic errors, while path length scaled the systematic errors logarithmically, similar to Weber-Fechner law. While we show that both models captured participant’s behavior, a model based on vector addition best captured the patterns of error in the homing vector. Our study therefore has important implications for how humans track their location, suggesting that vector-based models provide a reasonable and simple explanation for how we do so.

## Intro

“Dead reckoning,” first coined by Charles Darwin (Darwin, 1856/1987), described a process whereby experienced navigators kept course to a particular spot over long distances and changes in directions, despite being in the featureless arctic tundra. All animal species tested show the ability to dead reckon (referred to here as path integration), including spiders (Görner, 1958), bees (Lindauer, 1963), gerbils (Mittelstaedt & Mittelstaedt, 1980), hamsters (Etienne, 1987), house mice (Alyan & Jander, 1994), rats (Tolman, 1948), birds (Mittelstaedt & Mittelstaedt, 1982), ants (Wehner & Srinivasan, 1981), arthropods (Mittelstaedt, 1983), drosophila (Green et al., 2017), dogs (Seguinot, Cattet, & Benhamou, 1998), cats, and humans (Beritashvili, 1965). Please see Redish (1999), Gallistel (1990), and Klatzky, Loomis, and Golledge (1997) for a review of prior literature. Because humans employ vision as a primary modality to navigate, however, research on path integration has often been neglected in favor of situations in which visual input provides sufficient information to solve most navigational tasks, such as in desktop virtual reality. A limitation, however, with this testing modality is that it lacks the enriched cues and codes that we obtain when we freely move our body throughout space, thought to be critically important to path integration (Chance, Gaunet, Beall, & Loomis, 1998; Starrett & Ekstrom, 2018; Taube, Valerio, & Yoder, 2013).

Past experiments have often involved a path completion task in which the navigator is guided in physical space and must return using the shortest path back to the origin (Loomis et al., 1993; Görner, 1958; Müller & Wehner, 1988). Such work suggests that the navigator stores a representation of their current position relative to the origin that is periodically updated, frequently referred to as the homing vector. This in turn led to the suggestion that path integration involves vectorized representations of paths that are manipulated using vector addition, translation, rotation, and other well-established properties of matrix algebra. Computational modeling studies on path integration in both vertebrates and invertebrates support the idea of such vector-like representations, further suggesting that the homing vector is biased by systematic errors which are independent of random accumulated noise (Cartwright & Collett, 1987; Etienne et al., 1998; Etienne, Maurer, & Séguinot, 1996; Kubie & Fenton, 2009; Wittmann & Schwegler, 1995). Exactly how these pattern of systematic errors accumulate, however, is not clear, particularly in humans.

In humans, a frequently employed task is the triangle completion task in which the experimenter guides the participant on two legs of a triangle and then must return, without guidance, to the origin (Klatzky, 1990; Loomis, 1993). To model how systematic errors accumulate when human participants perform path completion tasks and the triangle completion task more specifically, Fujita et al. 1993 proposed the Encoding Error Model. This model proposes that the systematic errors in path completion tasks such as triangle completion task only occur during encoding stage. The model has four assumptions: (1) the internal representation satisfies Euclidean axioms (2) straight-line segments are encoded as a single value that represent their length (3) turns are encoded as a single value that represents the angle (4) there are no systematic errors during computation or execution of the homeward trajectory.

In support of their model, Fujita et al. fit data collected in Klatzky et al. 1990 and Loomis et al. 1993 involving the triangle completion task in the absence of vision. The model captured the systematic errors seen in both studies to a relatively high degree (see Fujita et al. 1993 Table 3). As predicted, though, the model performed poorly for paths with more than two sides or paths that crossed each other included in Klatzky et al. 1990. The Encoding Error model was expanded in Klatzky et al. 1999 to test its generalizability, who found that systematic errors were context and experience dependent. They also found that while partial vision increased path accuracy, it did not change the pattern of errors.

Another important finding, supported by the Encoding-Error Model and other studies (Petzschner & Glasauer, 2011), was that systematic errors in path integration, at least in small sized environments (≤10m), showed a pattern of regression to the mean. Specifically, past paths influenced the current paths and therefore, shorter angles and distances were overestimated and longer angles and distances were underestimated (Klatzky et al., 1990; Loomis et al., 1993). Petzschner and Glasauer 2011 (using desktop virtual reality) extended these findings by showing that the same angle or distance value could be overestimated in some cases and underestimated in others. The degree of under/overestimation depended on the distribution of priors, known as range effects, such that a broader distribution of priors (e.g., distances from 5-100 meters vs. 5-10 meters) increased the effect of the regression to the mean (Teghtsoonian & Teghtsoonian, 1978).

The issue of how the distribution of priors influences the current trajectories, however, begs the question of how path configurations affect errors in the triangle completion task. Specifically, past work suggests that the geometric properties of shapes can influence navigation (Cheng, 1986; Landau, Gleitman, & Spelke, 1981). For example, shapes like isosceles or equilateral triangles could serve as “templates” for how we learn paths (Seguinot et al., 1998) by providing a means for estimating paths that approximate it. Grid cells, neurons that fire as animals explore spatial environments, show 6-fold symmetry, with equilateral triangles composing part of this structure (Hafting, Fyhn, Molden, Moser, & Moser, 2005). Given arguments that neural codes might manifest in spatial representations useful for navigating (Bellmund, Gärdenfors, Moser, & Doeller, 2018; Milivojevic & Doeller, 2013) and the proposed link between path integration and grid cells (Chen, He, Kelly, Fiete, & McNamara, 2015; Moser & Moser, 2008), it could be the case that geometric regularities (equilateral triangles) also influence path integration. Indeed, some past studies on the triangle completion task provide support for the idea that geometric regularities can, in some cases, influence path accuracy (Klatzky et al., 1990). Yet, whether and how different types of triangles (equilateral vs. isosceles vs. scalene) influence path accuracy and patterns of errors on the unguided leg in the triangle completion task remains unclear.

Another important yet largely unanswered question about human path integration regards the accuracy and patterns of errors over longer distances. The vast majority of studies in human path integration have involved small-scale environments (<=10 meters) and consistent with this, computational models of path integration largely base their predictions on such smaller scales. For example, Klatzky et al., 1999 suggested that it is unlikly that same encoding function in their model is used for pathway that are larger than 10 meters^1^. A more recent computational model of path integration that employs grid cells suggests that, in the absence of specific mnemonic aids, path integration codes may rapidly degrade in mammals (Cheung, Ball, Milford, Wueth, & Wiles, 2012), consistent with the idea that path integration could breakdown dramatically over longer distances. Interestingly, however, other grid cell models assuming leaky integration rather than single value encoding suggest reliable estimations to up to 100 meters (Burak et al. 2009). Thus, an important question to test is how well human participants perform at path integration over longer distances (>=100 meters) and whether the Encoding-Error model vs. vector addition models most accurately captures such phenomenon in larger scales of space.

In the current study, we employed an omnidirectional treadmill and somatosensory input via handheld controllers (Figure 1A) to determine the extent to which manipulating the angle and distance participants needed to walk affected the accuracy of navigation without vision. The unique advantage of using the omnidirectional treadmill is it permits manipulation of infinity large spaces thereby eliminating the need for any boundaries while preserving the input from walking. The issue of boundaries, perceived or imagined, is a potential issue because if a participant were to over shoot a distance they would be stopped before hitting a wall, providing inadvertent feedback on the distances of the room and potentially affecting subsequent performance. In addition, the use of handheld controllers allowed us to carefully manipulate participant trajectories on the guided legs, an issue we return to in greater depth in the discussion.

**Figure 1.**
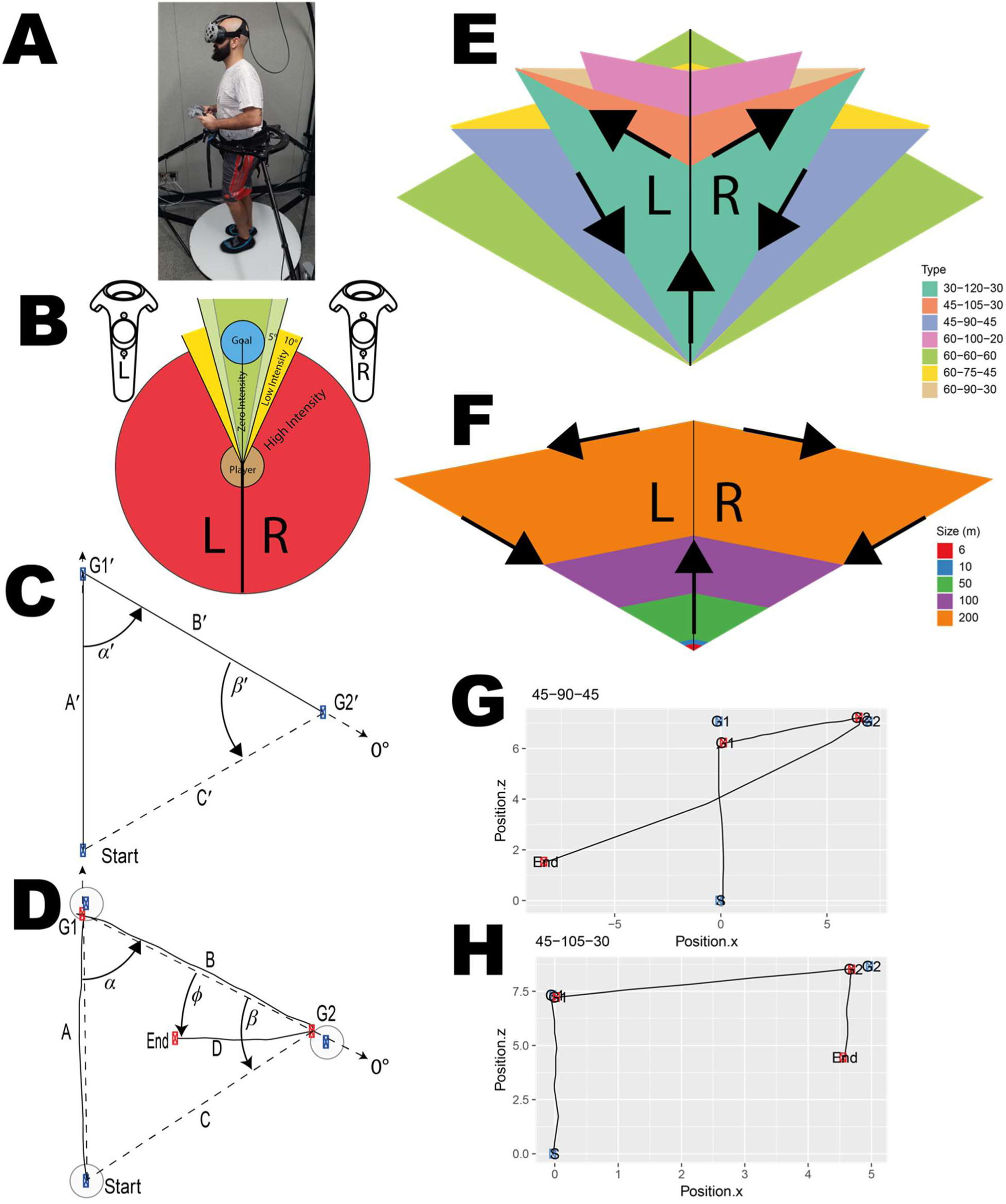
(A) HTC VIVE head set along with the handheld controllers using in the experiments, combined with Cyberith Virtualizer treadmill system to track participants in a much larger virtual environment while in stationary ambulation.(B)Visualization of HTC VIVE hand-held controllers feedback intensity based on the deviation of the angle. (C) Depiction of an equilateral triangle used in experiment 1. (D) Raw trace of participant’s path overlaid on the vector distances (dashed lines) between the points. The blue denotes the G1’ and G2’ locations that subject is being guided to and the red points are the subject’s unique G1 and G2 locations for that trial. (E) Triangle templets used in experiment 1 overlaid on top of each other and the legend denoting the internal angles. (F) Triangle templets used in experiment 1 overlaid on top of each other and the legend denoting the length of side C. (G) Raw trial where the participant over estimated distance and the angle. (H) Raw trial where the participant underestimated the distance and the angle

Here, we set out to test a simple yet novel model of path integration based on vector addition (often used to model path integration in other species) to better capture the pattern of errors in the triangle completion task in human participants (Etienne et al., 1998; Cartwright & Collett, 1987). Experiment 1 explicitly manipulated triangle type (while keeping homing distance constant) to test the extent to which different shapes of triangles (isosceles, equilateral) influenced how participants learned the homing vector. In Experiment 2, we explicitly manipulated the distance participants had to walk to reach the origin (while keeping triangle type constant) to determine how participants performed over a range of different distances. Critically, by manipulating these variables, we were able to simultaneously test hypotheses related to 1) triangle type and whether some might perform better than others; 2) homing distance and whether path integration would show different properties at ∼10m vs. ∼100m; 3) which model, one based on vector addition or the Encoding-Error model, would provide a better account of the pattern of findings. We provide a detailed comparison of the assumptions and set-up of the different models in the Methods section.

## Results

### Experiment 1

#### Basic behavior

An example raw trace of a participant’s path overlaid on the vector distances is shown in Figure 1D (dashed lines) between the points. We defined angle error as β - ϕ, where a positive number denotes an overshoot and negative an undershoot. Distance error is the ratio of leg D (unguided walked distance) over the distance of C (homing vector from G2); a value greater than 1 is an overshoot and less than 1 an undershoot. As can be seen in the raw example shown (Figure 1G & H) and others (Supplementary Figure 1), although participants were often quite accurate at completing the triangle, they tended to overestimate the angle and underestimate the distance, regardless of triangle type. We will compare our finding of systematic errors with prior literature, specifically, with Klatzky et al.,1999, in the Discussion section.

#### Participants overestimate angle and underestimate distance

We next addressed the extent to which this overestimation of angle and underestimation of distance was true across the group of participants. As shown in Figure 2A, we found a tendency for participants to overestimate the angle they needed to turn to reach their start point (t(21)=3.7,p<0.001, Cohen’s d=0.79,BF_10_>10), with participants, on average, tending to turn about 34.71°±9.37° too far when estimating the angle they would need to turn to reach the origin. In contrast, we found that participants tended to underestimate the distance they needed to walk to get back to the start point, with participants normalized walked distance significantly less than 1 (see Figure 2B, t(21)=16,p<0.001, Cohen’s d=3.42, BF_10_>10). Nonetheless, the average normalized walked distance was 0.87±0.05 (8.70m±0.50m), which was, on average, close to the correct response of 1 (10m). To determine the overall accuracy of the walked distance, we regressed the homing vector (leg C) onto participants’ unguided walked vector (leg D) using a vector model described in the methods. The beta values were positive and well above zero (t(21)=5.4,p<.001,Cohen’s d= 1.151, BF_10_>10), demonstrating that participants, despite underestimating distance, were well above chance in their estimates.

**Figure 2:**
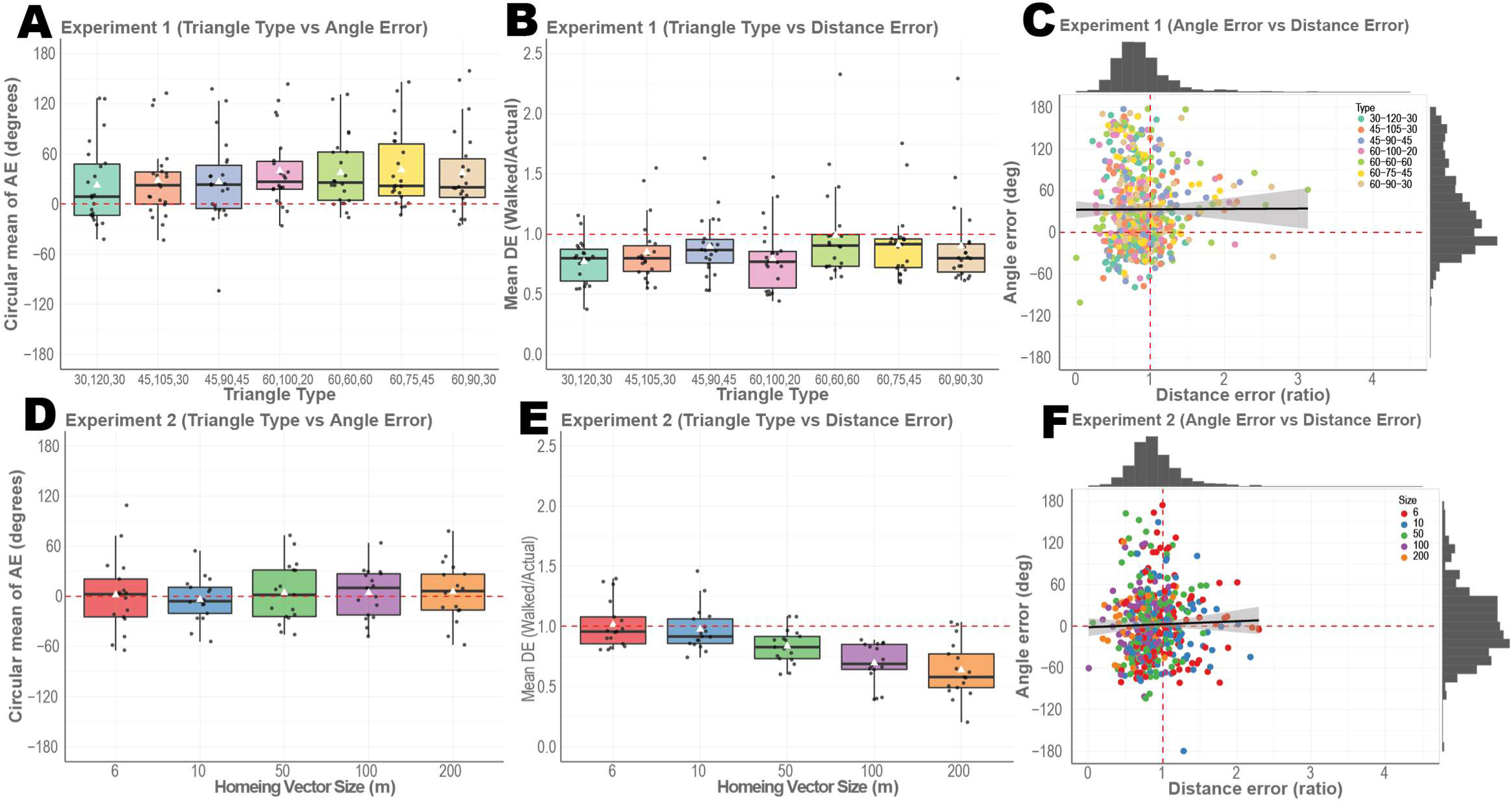
White triangles represent the mean while the median is shown as a black bar. (A)Circular mean of angle error for the 7 triangle types from experiment 1 (F(6,21)=2.9, p<0.01, η^2^ =0.058 BF_10_=1.72). (B)Mean distance error for the unguided walk from experiment 1 (F(6, 21)=2.1, p<0.1.33e-5, η^2^=0.109 BF_10_>10). (C) Angle error and Distance error of all trials from experiment 1 showing no correlation (t(579)=0.084,p=0.933, BF_01_>10). (D)Circular mean of angle error for the 5 triangle sizes (F(4,16)=0.609, p=0.658, η^2^ =0.036,BF_01_>10). (E) Mean distance error for the unguided walk from experiment 2 (F(4,16)=21.107, p<3.913e-11, η^2^ =0.553 and BF_10_>10). (F) Angle error and Distance error of all trials from experiment 2 showing no correlation (t(487)=0.623,p=0.533, BF_01_>7.8).

#### Results not dependent on the sensory modality of guidance information

To ensure that our results were not due to difficulty with employing the handheld controllers to navigate the guided legs, we compared against a subset of trials in Experiment 1 in which the guided legs involved a visual beacon (note that participants otherwise navigated the unguided legs identically in somatosensory and vision conditions). During the *guided* section of the trials, there was no effect of vision (Supplementary Figure 2A t(21)=1.09, p=0.288, Cohen’s d=0.336 and BF_01_>3), confirming that the hand-held controller feedback system provides sufficient guidance. For angle error on the unguided leg, as shown in Supplementary Figure 2B&D, we found a slight but significant improvement in the vision-on (SD:43.10°) compared to vision-off (SD:46.51°) condition (F(1, 21)=4.9, p<0.026, η2=0.016 BF_10_=1.16). For distance error, as shown in Supplementary Figure 2C&E, we also found a decrease in distance error during vision-on (SD:0.256) trials compared to vision-off (SD:0.271) (F(1, 21)=8.2, p<0.004, η^2^ =0.026 BF_10_=4). These findings suggest that providing vision on the guided legs did improve angle and distance estimates on the unguided leg, but that participants still tended to overestimate angle and underestimate distance (see Supplementary Figure 2D&H for additional information). Klatzky et al. 1999 also found partial vision to improve accuracy, though it seemed to have little effect the direction of systematic errors. Thus, the overestimation of angle and underestimation of distance that we observed cannot be accounted for by difficulty in completing the unguided legs using somatosensory input alone.

#### Little to no consistent effect of triangle shape on patterns of error in path integration

Next, we wished to address the issue of triangle shape and whether this may have contributed in any way to the patterns of errors for the unguided leg, as this might suggest participants used geometric features to anchor their path integration knowledge. For example, it could be that participants were most accurate for distance and angle on one triangle type (for example, right or equilateral triangles). To address this issue, we compared error on the unguided leg with triangle type as an independent factor. Overall, we found only a modest effect of triangle type on angle error (F(6,21)=2.9, p<0.01, η^2^ =0.058, BF_10_=1.72). Distance error, however, showed a fairly robust difference as a function of triangle type (F(6, 21)=5.7, p<0.1.33e-5, η^2^=0.109 BF_10_>10); see Figure 2A and 2B. Importantly, however, we did not find a consistent effect of triangle type across angle *and* distance errors, which might be expected if triangle *shape* had an influence on trajectories. For example, the isosceles triangle (30,120,30) showed the lowest mean angle error (10.96°±9.11°) yet the equilateral triangle demonstrated the lowest mean distance error (0.985±0.062). Thus, the inconsistent effects across triangle types and the small effects sizes we obtained for angle error suggest that participants were unlikely to be relying on geometric cues from the triangle shapes, which would involve remembering both the angle and distance for a specific triangle type. Instead, we attribute the lower angle and distance errors for isosceles and equilateral triangles, respectively, to the effects of repeating the same distances two times, an issue we return to in the Discussion.

As an additional analysis to investigate the use of geometric features of triangles, if participants were using specific shapes over others to perform the task, we might expect that both angle and distance errors would be correlated, consistent with using the shape, rather than individual features, to compute the unguided leg. Comparing angle and distance error is also important to determining the extent to which these two estimates were stored in a common vs. independent manner. We found no correlation between angle and distance error across trials and participants r(579)=0.0035, p=0.933 (Figure 2C), suggesting that angle and distance errors were not related to each other. We also observed no clustering of angle and distance error by triangle type (Figure 2C). Finally, we looked at the left and right handedness of the triangle and found no difference between them (Supplementary Figure 3A & B; angle error t(21)=0.7, p=0.485, Cohen’s d=0.118, BF_01_>3 and distance error t(21)=1.136, p=0.268, Cohen’s d=0.103 and BF_01_=2.53). Together, these findings suggest that triangle shape and the direction which participants navigated the triangle (i.e., right or left), contributed minimally, if at all, to performance on the unguided leg.

#### Computational modeling suggests that participants under and unevenly weigh the guided legs in Experiment 1

To better understand the pattern of errors that participants made in Experiment 1, we built a computational model to predict the pattern of errors for the unguided legs. We combined angle and distance into a single vector value (see Methods) and employed the vectors for guided leg A and B as predictors for the unguided leg. Based on previous findings (Fujita et al., 1993), we would expect the guided legs to strongly predict performance on the unguided leg. The modeling approach we employed also allowed us to compare the relative weighting of leg A vs. leg B and whether past trial history had any impact on unguided leg performance.

The modeling analysis revealed that both guided legs A and B strongly predicted performance on the unguided leg (mean *β_A_* = 0.3, t(21)=2.86 p<0.0001 BF_10_>3 and mean *β_B_* =0.813, t(21)=7.41 p<0.0001, BF_10_>10; Figure 3A). Notably, only A’s beta value was significantly less than 1 (t(21)=6.6, p<1.299e-6, BF_10_>10), suggesting that participants underweighted leg A when estimating the return vector, potentially, accounting for the angle overestimation. In addition, leg B was weighted higher than leg A, (t(21)=3.62, p<0.002, Cohen’s d=1.02, BF_10_>10).

**Figure 3.**
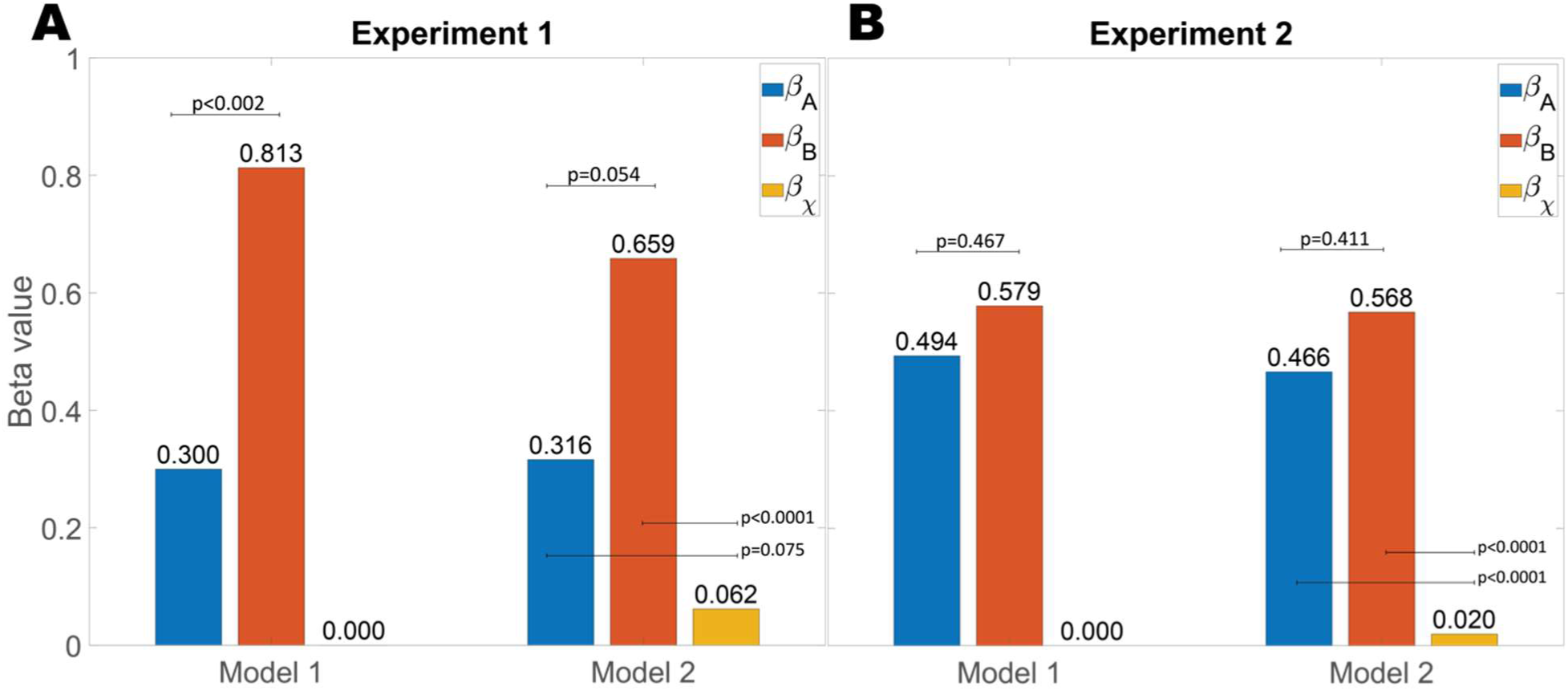
Mean Beta values from the vector model for (A): experiment 1 and (B): experiment 2.

For model 2 (equation 5), which included participants’ past trial history, we found mean *β_A_* = 0.316, t(21)=2.52, p<0.02, BF_10_=2.75 and mean *β_B_* =0.659, t(21)=5.51, p<0.0001, BF_10_>10, suggesting similar results in terms of underweighting the guided legs as Model 1. However, we found no significant effect of past trials (mean *β_x_* =0.062, t(21)=1.03, p<0.31, BF_01_=2.89), suggesting that sequential effects were minimal in Experiment 1 (Figure 3A). Because the priors were relatively stable in Experiment 1 (i.e., distance was not explicitly manipulated), this result is consistent with the idea that the range of distances tested in Experiment 1 was insufficient to see a regression of to the mean effect (Teghtsoonian & Teghtsoonian, 1978).

Taken together, these findings suggest that the patterns from Experiment 1, which involved different triangle types, could be captured by our vector-based models, particularly Model 1. Participants underweighted both guided legs A and B, with a tendency to underweight leg A to a greater extent. We found no evidence for distances and angles on past trials providing any explanatory power for the unguided leg.

#### Model validation

Next, we simulated Model 1 to determine whether it could account for the trends observed in the empirical data (Palminteri, Wyart, & Koechlin, 2017; Wilson & Collins, 2019). We found that Model 1 captured both the angle overestimation (Figure 4A) and distance underestimation (Figure 4B) in Experiment 1. The simulation results also supported the idea that Model 1 provided a better account for the data than Model 2 (Figure 5A-C) and captured the relevant empirical phenomenon reported here.

**Figure 4:**
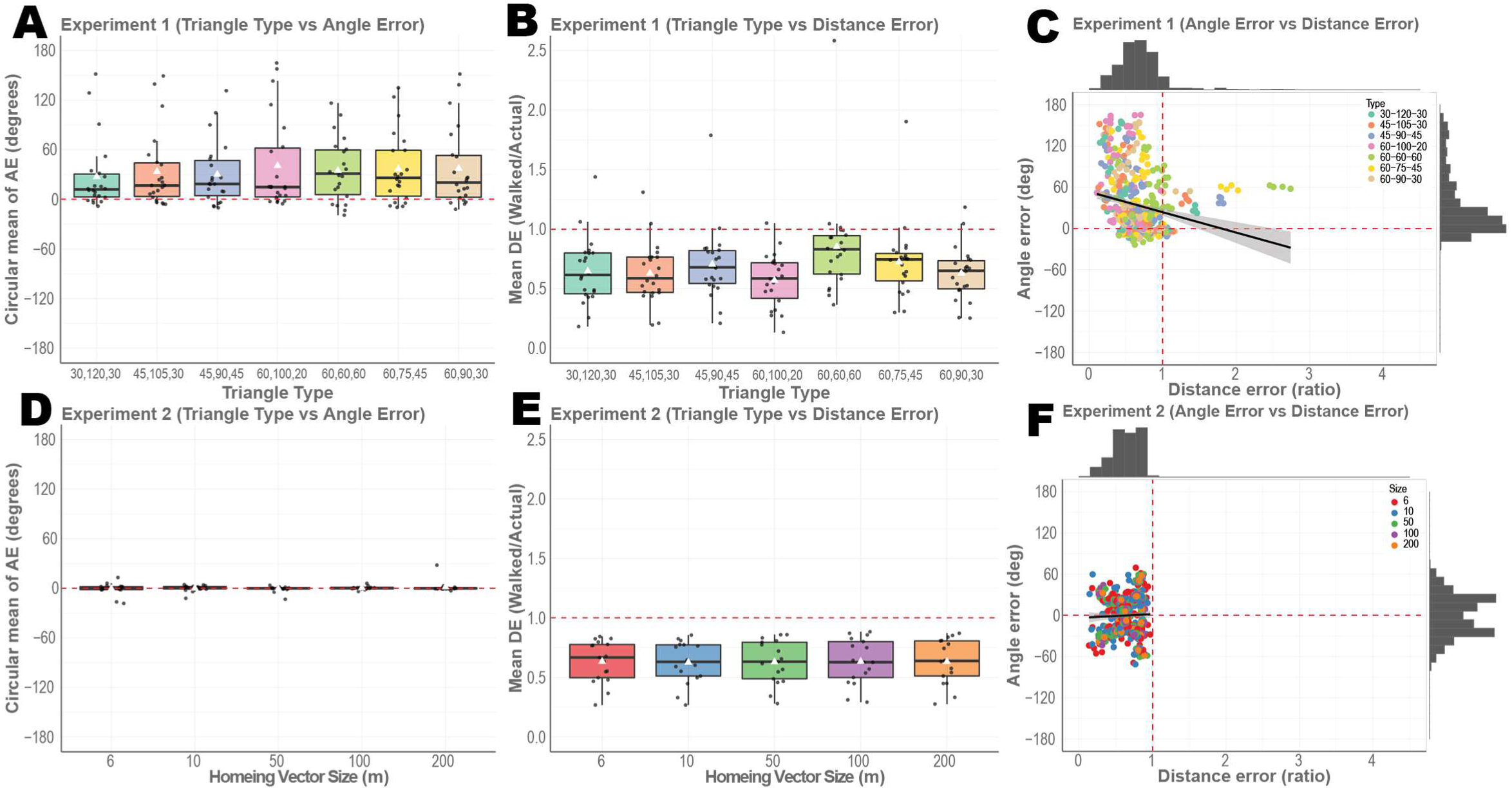
White triangles represent the mean while the median is shown as a black bar. Simulated data from Model 1 (A)Circular mean of angle error for the 7 triangle types from experiment 1. (B)Mean distance error for the unguided walk from experiment 1. (C) Angle error and Distance error of all trials from experiment 1 showing no correlation. (D)Circular mean of angle error for the 5 triangle sizes. (E) Mean distance error for the unguided walk from experiment 2. (F) Angle error and Distance error of all trials from experiment 2 showing no correlation.

**Figure 5:**
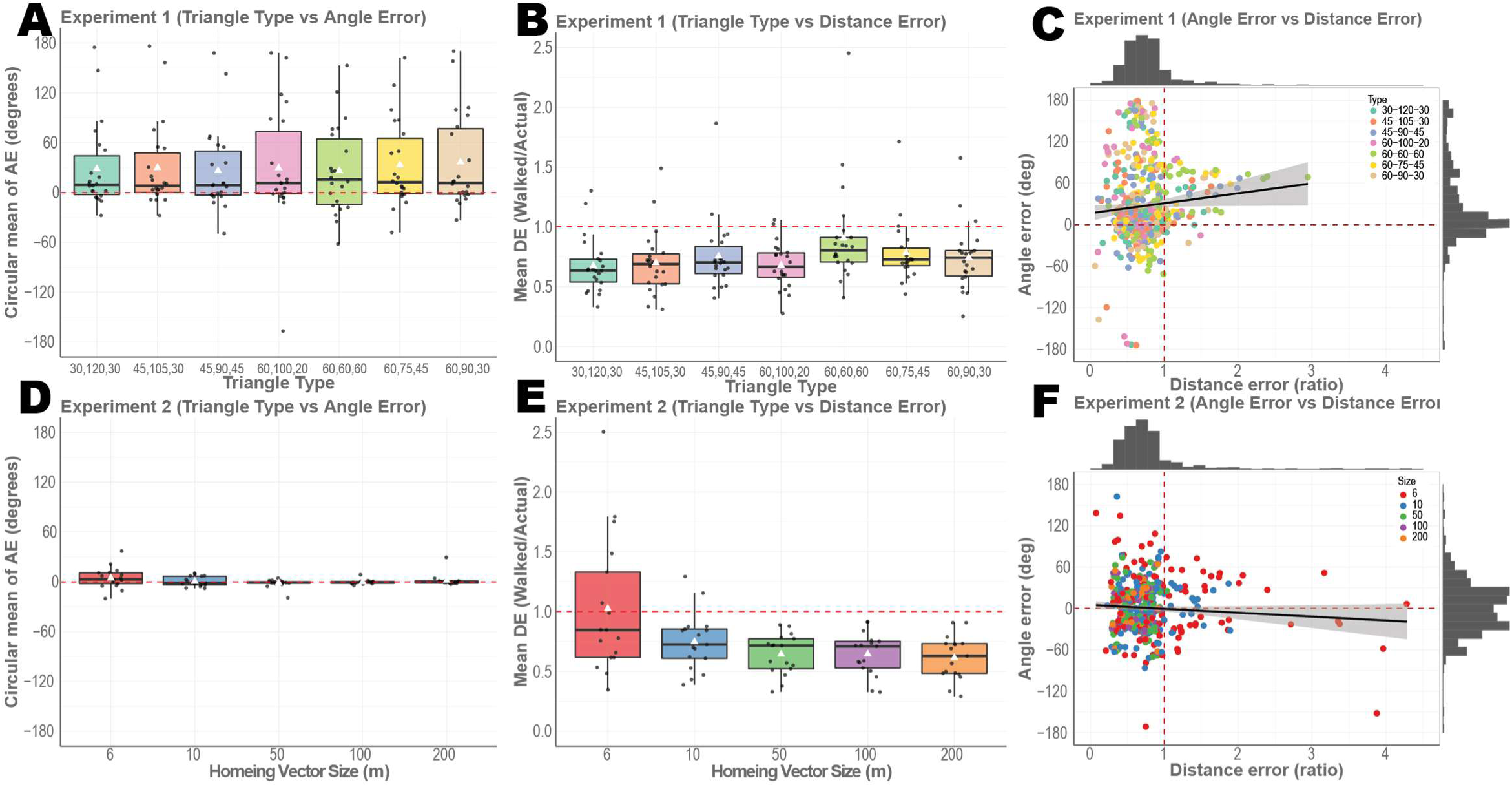
White triangles represent the mean while the median is shown as a black bar. Simulated data from Model 2 (A)Circular mean of angle error for the 7 triangle types from experiment 1. (B)Mean distance error for the unguided walk from experiment 1. (C) Angle error and Distance error of all trials from experiment 1 showing no correlation. (D)Circular mean of angle error for the 5 triangle sizes. (E) Mean distance error for the unguided walk from experiment 2. (F) Angle error and Distance error of all trials from experiment 2 showing no correlation.

#### Encoding-Error Model

We fitted and simulated our data using the Encoding-Error Model, and, similar to Model 1 and Model 2, were able to capture the systematic errors in angle overestimation (Figure 6a) and distance underestimation (Figure 6b). Similarly, the Encoding-Error Model, given the limited range of triangle distances in Experiment 1, did not show regression to the mean. When we directly compared the models (Supplementary Figure 6 A-C), however, we found that Model 1 fit the data fairly decisively, at both subject and group level. While Model 1 did outperform the other two models in BIC and AIC, the confusion matrix in Supplementary figure 7 A-C showed that simulated data from Encoding-Error model did not fit Encoding-Error model best compared to the two vector addition models. This method of model recovery suggests some limitations with our model comparison (i.e. how well our task can distinguish between models) and was likely due to small number of trials and the fact that the vector addition models involved far fewer free parameters than the Encoding-Error Model (Wilson & Collins 2019). We return to a more detailed comparison between vector addition and Encoding-Error Models in the Discussion.

**Figure 6:**
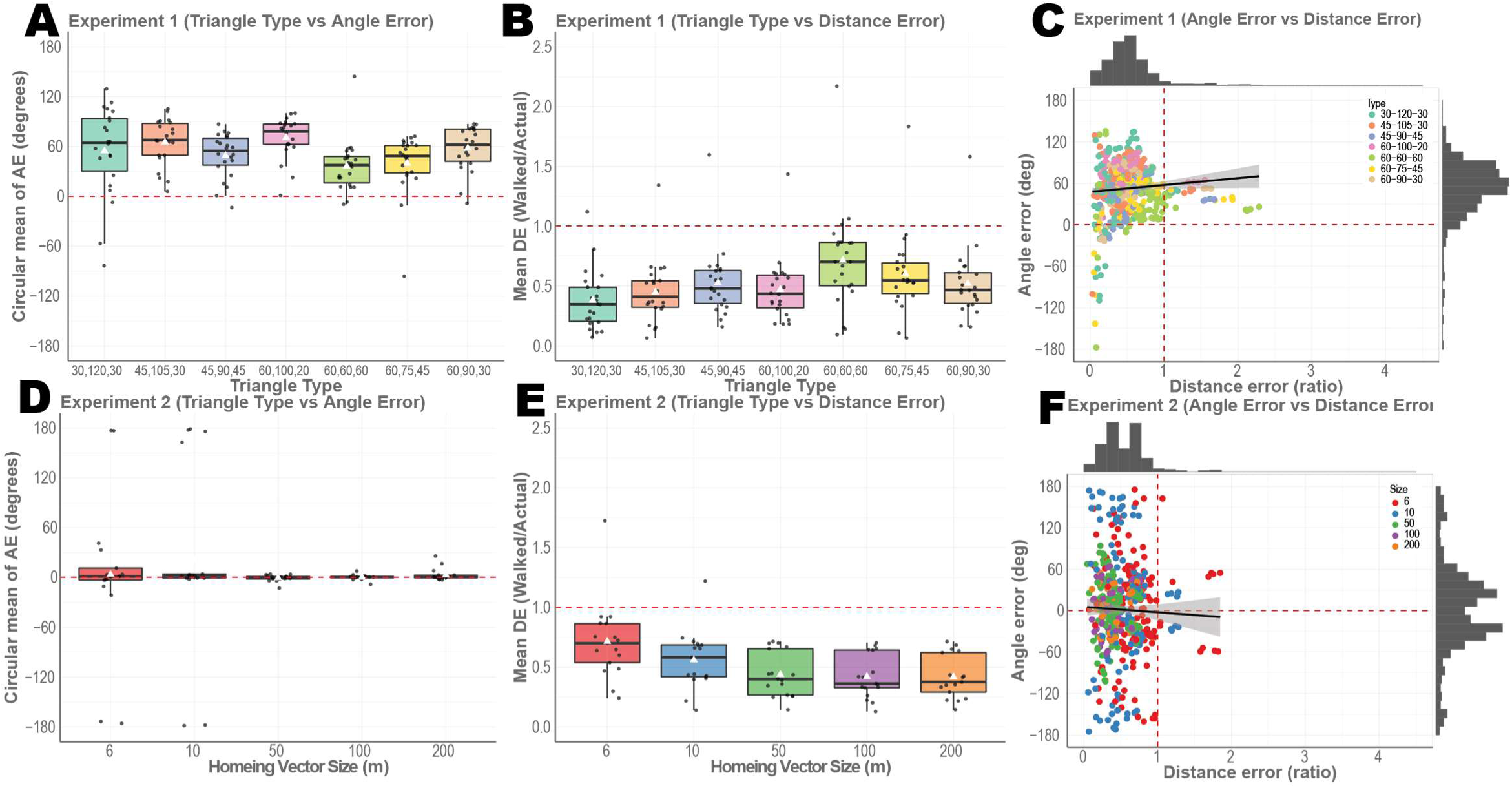
White triangles represent the mean while the median is shown as a black bar. Simulated data from Encoding-Error Model (A)Circular mean of angle error for the 7 triangle types from experiment 1. (B)Mean distance error for the unguided walk from experiment 1. (C) Angle error and Distance error of all trials from experiment 1 showing no correlation. (D)Circular mean of angle error for the 5 triangle sizes. (E) Mean distance error for the unguided walk from experiment 2. (F) Angle error and Distance error of all trials from experiment 2 showing no correlation.

**Figure 7:**
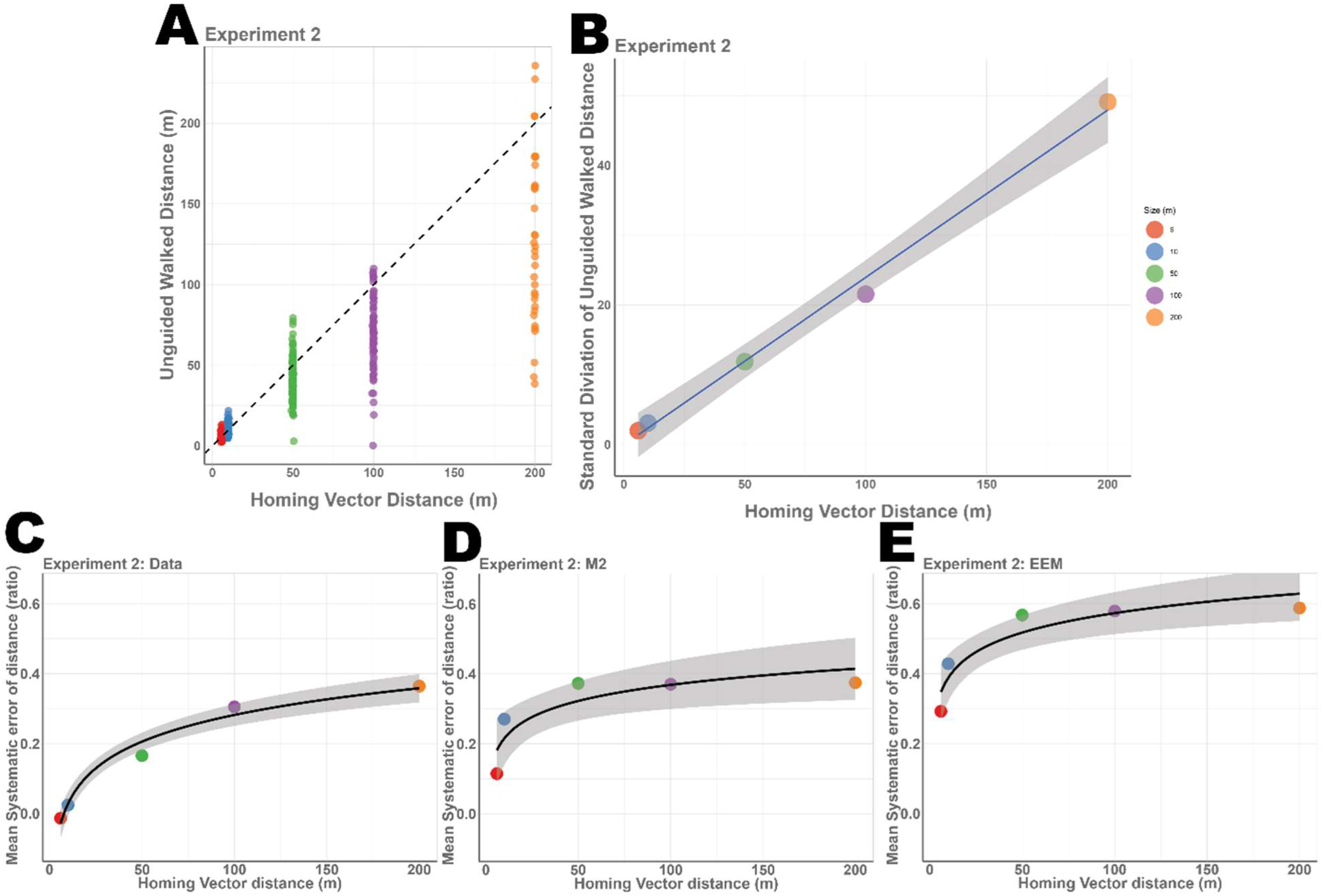
Results from Experiment 2, (A)showing the distribution of the Unguided walked distances for each triangle size with y=x plotted at the dotted line. (B) Standard deviation of the Unguided walked distances show a linear increase (t(4)=23.6, p<0.0001) (C) showing mean systematic errors of distance (1- distance error) increases logarithmically (t(4)=11.65, p<0.001). (D)showing mean systematic errors of distance of the simulated data from model 2 increasing logarithmically (t(4)=3.187, p<0.05). (E)showing mean systematic errors of distance of the simulated data from Encoding-Error Model increasing logarithmically (t(4)=4.407, p<0.022).

### Experiment 2

#### Basic behavior

In Experiment 2, we manipulated the distance of the triangles (perimeters = 15.19, 25.32, 126.60, 253.20, and 506.42 meters) while keeping triangle geometry relatively constant. This involved necessarily manipulating the distance of the guided legs, yet we overall maintained a scalene triangle shape, thus leaving angle as comparatively constant as possible. We implemented the same task structure as Experiment 1 but here we kept the shape of the guided path the same and varied the scale across trials.

#### Participants systematically underestimated distance but accurately estimated angle

For angle error, somewhat in contrast to Experiment 1, we found no significant overestimation or underestimation of angle, with participant’s showing a mean error of 0.8°±7.44° (t(16)=0.107, p=0.916, Cohen’s d=0.026, BF_01_>3). We also found no effect of triangle size on angle error (Figure 2D (F(4,16)=0.609, p=0.658, η^2^ =0.036,BF_01_>10). We attribute this to the fact that triangle configuration was consistent across Experiment 2, as we primarily manipulated distance.

We found evidence of fairly accurate estimation of distance for smaller triangle perimeters (15-25m perimeter) and considerable underestimation for larger triangle perimeters (126m – 500m perimeter). In fact, we found a trend whereby distance underestimation increased as a function of the unguided distance (Figure 2E, F(4,16)=21.107, p<3.913e-11, η^2^ =0.553 and BF_10_>10). This is shown in Figure 7A, where the dotted line indicates a slope of 1, with the actual slope well below this value. In other words, the further that participants walked, the more they tended to underestimate the unguided leg.

To better understand this phenomenon, we analyzed the spread of the errors as participants walked the unguided leg. We found that distribution of distance error scaled linearly as a function of the walked distance. As shown in Figure 7B, the standard deviation of the walked unguided distances increased linearly, as shown by a regression fit (F(1,3)=557.4, *β*_1_ =4.417,r^2^=0.9929), suggesting that the greater the walked distance, the proportionately greater the error in distance with variance increasing exponentially. Note that this phenomenon is distinct from that related to systematic error. Systematic errors for distance error increased as well, however, this increase was best fit by a logarithmic function (Figure 7C, t(4)=11.65, p<0.00136) rather than linearly, similar to Weber–Fechner and Stevens’ power law (Stevens,1975). Together, these findings suggest that as participants walked longer distances, they tended to increase their underestimation of the distance they would need to walk and scale their errors logarithmically as a function of distance.

Similar to Experiment 1, we also found no correlation between angle and distance error (Figure 2C, t(487)=0.623,p=0.533, BF_01_>7.8). We also found no effect of right vs. left turns on guided legs (angle error: t(16)=1.51, p=0.151, Cohen’s d=0.245, BF_01_=1.55 and distance error: (t(16)=0.724, p=0.4797, Cohen’s d=0.176 and BF_01_=3.188), see supplementary Figure 3C & D.

#### Computational modeling suggests sequential effects of past trials in Experiment 2

To better understand the effects of the guided legs on the unguided leg estimates in Experiment 2, we employed the computational model used in Experiment 1 to predict the pattern of errors for the unguided legs. The modeling analysis again revealed that both guided legs A and B strongly predicted performance on the unguided leg (mean *β_A_*=0. 494, t(16)=5.09, p<0.0001, BF_10_>10 and mean *β_B_*=0. 579, t(16)=9.29, p<0.0001, BF_10_>10) (Figure 3B). Notably, both beta values were less than 1 (*β_A_* t(16)=5.22 p<2.24e-5, BF_10_>10 and *β_b_* t(16)=7.07 p<0.0094, BF_10_>10), suggesting that participants underweighted *both* legs when estimating the return vector. In addition, unlike Experiment 1, both legs were weighted evenly (t(16)=0.63,p=0.467, Cohen’s d=0.25, BF_01_>3). These findings are perhaps unsurprising because angle was neither under nor overestimated.

Comparing model 1 (modeling the distance of the guided legs to predict the unguided legs) and 2 (using model 1 with an additional term for past trial distances), we found significant fits for all three beta terms. In other words, guided legs A & B, as well as past trial history (mean *β_A_* = 0.466, t(16)=4.42, p<0.001, BF_10_>10, mean *β_B_* =0.568, t(16)=9.29, p<0.001, BF_10_>10 and mean *β_x_* =0.020, t(16)=3.82, p<0.001, BF_10_>10), all predicted errors in walking the unguided leg in Experiment 2. Thus, in contrast to Experiment 1, trial history provided a significant explanation of error in Experiment 2.

#### Model validation

Next, we simulated our data in a manner similar to Experiment 1. Simulated data from Model 1 showed that we were able to capture participant patterns in angle error (Figure 4D). While Model 1 captured the distance underestimation (Figure 4E), it did not capture the trend of increase in underestimation as a function of distance. We hypothesized that this effect could be an influence of past trials, in other words, a form of regression to the mean (Klatzky, Beall, Loomis, Golledge, & Philbeck, 1999; Petzschner & Glasauer, 2011). Figure 5D shows the simulated angle error from Model 2, and we are again able to capture the accurate angle predictions. Importantly, however, simulated distance error, as shown in Figure 5E, better captured the pattern of distance underestimation. Model 2, in particular, captured the tendency of participant underestimation of distance to increase as a function of distance while Model 1 (which did not include trial history) was not able to capture this effect. These findings suggest that the increasing underestimation of distance was influenced, in part, by past trials.

#### Encoding-Error Model

The Encoding-Error Model also captured some of the same patterns in the data as Model 1 and 2. The simulated data from the Encoding-Error Model showed accurate angle error and underestimation of distance errors as a function of distance (Figure 6 D&E). We also considered how well the Encoding-Error Model compared with Model 2 in terms of capturing the mean systematic error in distance, which was 1-mean distance error (Figure 7 C-E). While the Encoding-Error Model fit the logarithmic function of systematic errors, the values were less accurate than Model 2. Similar to Experiment 1, Model 2 best fit the data but the BIC and AIC favored Model 1 (Supplementary figure 6 D-F). Notably, though, our analyses (see Figure 4 D-F) suggested that Model 1 did not capture the pattern of systematic errors and thus we removed it from the model comparison with the Encoding-Error Model. As shown in Supplementary Figure 8 A-D, we can see Model 2 fits 11 subject’s data better while Encoding-Error Model fit the other 5 subject data better. Similar to Experiment 1, the confusion matrix (Supplementary Figure 9 A-C) showed that Encoding-Error model did not fit its own simulated data well. This was likely due to small number of trials and the fact that the vector addition models involved fewer free parameters than the Encoding-Error Model (Appendix A). We return to a more detailed comparison of the models in the Discussion.

## Discussion

In two different experiments, participants were guided on two legs of a triangle and then attempted to return to the origin without any input using a novel interface involving an omnidirectional treadmill. In Experiment 1, we manipulated triangle type (equilateral vs. isosceles vs. right vs. scalene) while holding distance on the unguided leg constant to minimize prior effects. Consistent with previous work using the triangle completion task in small-scale room sized environments (Fujita et al., 1993; Klatzky et al., 1997; Loomis et al., 1993; Philbeck et al., 2001; Yamamoto et al., 2014), we found that participants underestimated distance and overestimated angle, however these systematic errors did not show a regression to the mean effect. In Experiment 1, our computational modeling results suggested that this pattern could be explained by a model in which participants underweighted leg A compared to leg B. In Experiment 2, we found systematic errors in distance as participants accurately estimated the angle they needed to turn while increasingly underestimating the unguided leg as a function of distance, consistent with logarithmic scaling described in the Weber-Fechner law. Modeling results for Experiment 2 further suggested equal weighting of both encoded legs. We also found no correlation between angle and distance errors in both experiments, consistent with reports that, at least in part, we derive angular motion from the semicircular canals and linear motion through the otoliths (Carriot et al., 2015). Our findings thus suggest that participants used independent estimates of direction and magnitude to estimate a homing vector, with the current trial guided legs influencing estimates of the homing vector.

In Experiment 1, we found that triangle type had little influence on participants’ performance on the unguided leg. For example, it might be possible to predict that equilateral triangles or right triangles would be overall more accurate than scalene triangles. This is because these geometries are far more regular and potentially easier to encode holistically, particularly given their influence on visually guided navigation (Moar & Bower, 1983). While we did find that the equilateral triangle showed significantly lower distance error, we attribute this to a working memory effect based on the equivalence of all three leg distances. Similarly, we found a weak tendency for the isosceles triangle angles (30,120,30) to show lower angle overestimation. One might expect, though, that if participants, or a subset of them, used a template (i.e., fit the guided legs to an equilateral or right triangle and estimated the return vector from there), we would also find that they would also be more accurate on that particular triangle. While we did not find this over our group of participants, we did find two participants who showed a high degree of accuracy on equilateral and right isosceles triangles (Supplementary Figure 4C). Thus, while it is possible a small subset of a participants employed triangle “templates,” our findings suggest that the majority of participants did not. Overall, the lack of any consistent effects in angle and/or distance for specific triangle types in terms of accuracy and the lack of a correlation between angle and distance representations suggests that the geometric properties of specific triangles played little, if any, role in solving the triangle completion task. Instead, these findings thus suggest that path integration mechanisms in humans are based on continuous encoding of heading direction and magnitude during the guided legs, after which vector addition is used to construct a homing vector.

In Experiment 2, we tested homing behavior over distances much longer than those typically employed in past human studies. Almost all of our current knowledge base about path integration mechanisms in humans derive from testing in room-sized environments, and therefore, in contrast to what is known about other species, the extent to which path integration mechanisms operate accurately over distances greater than 10 meters remains unclear. We found that participants were fairly accurate in their ability to complete the third leg of a triangle, even for triangle perimeters as long as 500 meters. Although we found a systematic increase in error and underestimation as a function of longer distances, these biases increased logarithmically, suggesting that the basic mechanisms underlying path integration were not substantially different at 500 meters compared to 25 meters. In contrast to Experiment 1, we found that both legs A and B contributed equally to errors in unguided leg C, although we attribute this effect to the fact that we did not manipulate angle in Experiment 2. We did find, however, that past trial history contributed significantly to the pattern of errors at longer distances. These findings suggest that in fact some of the properties of path integration do change somewhat over longer distances, particularly the tendency to erroneously weight past trials to estimate the current ones. Given that our two models, however, involved the same basic conceptual set-up (leg A+B=C), these results suggest that the basic mechanism of adding vector values for the guided legs to compute a homing vector held constant across experiments.

In one previous study, participants were blindfolded and attempted to walk in a straight line for several hundred meters in a desert environment. In contrast to our findings, this study found that participants tended to walk in circles, even as early as 10 meters into their 1 kilometer leg (Souman et al., 2009). This, in turn, might suggest that path integration mechanisms in humans undergo a form of catastrophic breakdown at longer distances. There are several key differences between our study and that of Souman et al., however. Perhaps most importantly, our study involved participants encoding distances and angles they had turned to estimate a new vector back to the origin. Goal directed navigation involves fundamental differences from simply walking in a straight line (Klatzky et al., 1997), and it is possible that having a specific goal location in our task (return to the origin) reduced the tendency to walk in circles. Another important difference between our studies is that participants navigated on an omnidirectional treadmill while in the Souman et al. study, they navigated in the real-world (desert environment). Could our treadmill have prevented participants from taking circuitous paths? We analyzed all paths in the treadmill and did indeed find some examples of circuitous paths, suggesting that the treadmill interface itself did not preclude participants from employing this (Supplementary Figure 5). We also note that another study from the same group employed an omnidirectional treadmill interface, finding that participants walked in largely comparable ways to how they might in the real world (Souman et al., 2011). While we cannot rule out other differences between our experiments, we note that we found similar results for the triangle completion task in small-scale space as previously reported in room-sized environments, and thus we believe that the interface itself is unlikely to account for the differences in our findings. Instead, we favor an account based on the importance of using path integration mechanisms to find the origin.

Our computational modeling results indicated an effect of past trials on participant error patterns in Experiment 2 but not Experiment 1. In other words, for the longer distance triangles, we found a weak, but significant bias for past trials to influence the extent to which participants underestimated the amount they needed to walk on the current triangle. For large triangles, therefore, shorter past trials would result in a greater tendency to underestimate distance. Notably, including the history term in our model significantly increased our ability to account for the increasing tendency of participants to undershoot the distance they needed to walk on the unguided leg. These findings support the idea that for particularly long distances, path integration is also influenced by a form of regression to the mean from past trials, thus explaining why undershoot increased with longer distances. These findings, which, to the best of our knowledge, have not been demonstrated previously at such long distances in humans, suggest that path integration is not merely a function of the current walked triangle, but is also influenced by the memories and experiences of past trajectories.

Because of our strong reliance on visual input, testing humans in the absence of vision is challenging, particular due the possibility of trip hazards and collisions. Thus, many researchers have chosen to investigate path integration using desktop VR, which also allows simultaneous brain imaging, for example, using fMRI (Chadwick, Jolly, Amos, Hassabis, & Spiers, 2015; Chrastil, Sherrill, Hasselmo, & Stern, 2015). One limitation with desktop VR, however, is that it lacks the rich cues that one obtains from freely moving the body in space (Starrett & Ekstrom, 2018). These include vestibular information from head turns, proprioceptive information about body position, efferent copy from motor movements, and somatosensory input from the feet as they move over the surface (Gallistel, 1990; Lackner & DiZio, 2005; Loomis & Beall, 1998; Matthis, Yates, & Hayhoe, 2018; Visell, Giordano, Millet, & Cooperstock, 2011; Waller, Loomis, & Haun, 2004). Our novel interface was able to reproduce many of these cues, particularly those that would be expected from turning and shuffling the legs and feet. As such, we were able to capture novel aspects about non-visual navigation otherwise difficult to observe. Additionally, participants in our study generated their linear and angular motion, while non-VR versions of the triangle completion task used in the past relied on the experimenter physically guiding the participant’s movements. Previous versions of path completion task have used an object (rod or rope) in which the experimenter guides the participants by pulling or lowering for turning (Klatzky et al., 1990, Loomis et al. 1993, Klatzky et al., 1999). In contrast, in our design, participants received feedback from hand-held controllers indicating which way to go. We believe that the use of feedback via handheld controllers, rather than external forces to guide subjects, better approximates active walking. Specifically, active walking requires one to initiate the movement while outside forces that initiate or guide the movement would typically be referred to as passive. We believe by controlling for active walking during the guided portion, we have better controlled for differences between guided and unguided conditions. While the distinction between active and passive movement is a subtle one, recent work suggests important differences between these two forms of walking in terms of their neural bases (Carriot, Jamali, Brooks, & Cullen, 2015).

### Model comparisons: Vector addition models more plausible then Encoding-Error Model

Vector addition has long been assumed to be the functioning principle for path integration (Cartwright & Collett, 1987; Etienne et al., 1998; Kubie & Fenton, 2009). The vector addition models proposed in this paper (Models 1&2) assume that the homing vector is updated by summing vector representations of legs A and B. In contrast, the Encoding-Error Model assumes that the homing vector is created using the distance and angle values experienced during the entire guided portion. While both models are similar in aim, we believe the computational principles for the vector model may be more plausible. To employ the Encoding-Error Model, participants must form a representation of the linear relationship between distance guided and distance walked (distance representation) as well as for turns, for each path configuration across all subjects and trials. In addition, it is not clear whether the parameters of these linear functions generalize across studies and participants (Klatzky 1999). In contrast the Vector Addition Models assume a linear relationship between the guided leg and encoded vector, with the possibility of prior encoded vector values influencing the current trajectory.

As mentioned in the introduction, there are other reasons to think that vector addition models confer advantages, particularly in accounting for human path integration findings from the triangle completion task. The Encoding-Error Model has four requirements, with one important assumption being that the internal representations must obey Euclidean axioms. Recent papers, however, suggest that human spatial navigation, in some instances, may be better characterized by representations based on non-Euclidean labeled graphs (Warren, 2019). Specifically, Warren et al 2019 described path integration using simple vector manipulations with such manipulations preserved in non-Euclidean spaces. Our model, which can be readily adapted to non-Euclidean geometries, would therefore also provide greater flexibility than the Encoding-Error Model in terms of fitting violations of Euclidean axioms.

Another requirement of the Encoding-Error Model is the assumption that all systematic errors occur during encoding rather than during spatial reasoning or execution. Vector Addition Models are more flexible, assuming systematic errors can aggregate at different stages, whether it is during encoding, retrieval or computation of homing vector.

The Encoding Error Model, however, is limited in that leg A and B derive from the same linear function, such that leg A cannot be underestimated more than leg B. There may be instances, however, in which a leg is weighted differently in a path with 2 segments compared to 5 segments (Wan, Wang, & Crowell, 2013). In addition, the Encoding-Error Model is limited to 2 segmented triangular paths, based on the law of cosines (Appendix A), and does not perform well with 3 segmented paths (Fujita, Klatzky, Loomis, & Golledge, 1993). In contrast, vector addition models can readily be extended to n paths with the caveat of adding a free parameter with each segment. Notably, the vector addition models we employed here provided an overall better fit of the actual data (Supplementary figure 6 A&D), however the Encoding-Error model cannot be fully distinguished during model recovery (Supplementary figure 7). The likely reason for this is the small number of trials our task. While both Model 2 and Encoding-Error model can account for some patterns in the data, including systematic errors, importantly, Model 2 has the best log-likelihood fit (supplementary figure 6 A & D), despite the Encoding-Error model having more free-parameter. Overall, therefore, we think the vector addition models provide a better fit of our data and are parsimonious although more work is needed to allow a detailed and formal model comparison.

#### Limitations of the Vector Model

While the vector addition models employed here do a fairly good job of capturing the patterns of our findings in the two experiments in this study, they are not without certain limitations. One issue is that the model in its current form assumes that Leg A and Leg B are encoded with similar directions (i.e. 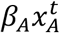 has the same direction as 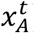) or opposite directions and only the vector magnitudes affect systematic errors. We hope to address the issue of vector directions in more detail in future models.

## Methods

### Training and the triangle completion task

All studies were approved by the UC Davis Institutional Review Board (IRB) with participants in some cases receiving class credit for their involvement. We employed a task used previously to investigate human path integration termed the triangle completion task (e.g., Loomis et al., 1993). Briefly, the task involves guiding participants on two legs of a triangle and then completing the third leg without guidance or feedback. Based on our goal of studying a variety of different triangle types and sizes, we adapted the task to an omnidirectional treadmill, the Cyberith Virtualizer treadmill. The task involved participants walking on the treadmill, with guidance on two of the legs provided by somatosensory feedback from HTC VIVE hand-held controllers. Participants wore the HTC VIVE headmounted headset to allow us to track head and body position, as well as to limit visual input.

To first ensure that participants could walk comfortably in the treadmill, we employed a pre-experimental training session. We employed an HTC VIVE head-mounted display to give visual feedback to ensure balance and comfort on the treadmill. In the first part of the training, we included a 3-stage puzzle game created in Unity 2017.1.1f1 in which participants had to explore an environment to find an object. Once participants completed the 3-stage puzzle game, reported no cybersickness, and the experimenter determined that their walking technique was adequate, they advanced to the next level. At this point, we introduced the HTC VIVE hand-held controllers feedback system (Figure 1B) and had subjects walk straight lines with no visual information while receiving feedback from the hand-held controllers. This insured that they could accurately perform the guided legs. Following this, they performed a small number of practice triangles. After practicing the triangle completion task on 6 unique triangles, which were not included in the experiment, the experiment started. The training period ranged from 30-60 min. To ensure participant safety, we occasionally questioned them about how they were feeling to guard against issues with cybersickness.

Participants then proceeded to the main experiment. The first experiment involved manipulating triangle geometry (i.e., primarily the angles they turned) and Experiment 2 involved manipulating triangle size (i.e., we manipulated the distance they walked on the third / unguided leg). Trial sequences were randomly chosen from 5 pseudorandomized configurations. In both experiments, we guided participants along the first two legs of the triangle using the hand-held controller feedback system (Figure 1B). The feedback system was designed such that if the participants strayed from their path, the controller vibrated accordingly to help guide them in walking in a straight line. When participants walked in the correct direction, the controller did not send feedback, allowing for active walking (passive guidance). Participants were guided along leg A’ and then along leg B’ by controller feedback (Figure 1C). At G2’, the hand-held controller feedback system turned off and participants were instructed to find their way to the start point. Participants pressed the trigger on the handheld controllers once they believed that they reached the start point. We constructed trial specific vectors to capture the performance variability during guided legs (see Figure 1E). We manually inspected these trials, and those which showed a clear deviation from linearity were excluded, which resulted in approximately 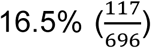 of removal of trials from Experiment 1 and 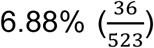 from Experiment 2 across participants. Participant data that exceeded 25% removed trials were excluded from the analysis. We redid the analysis by including all trials and participants and obtained similar results to what are reported here.

## Modeling

### Description of models

To further understand how the guided legs contributed to the angle and distance errors of the unguided leg, we created a vector model of path integration. In this model, we assume that participants estimate a “homing vector”, 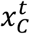, by combining the vectors corresponding to each of the guided legs for that trial (t), 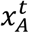 and 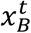. If path integration were optimal, people would combine these vectors in the following way

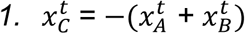

and would return perfectly to the point of origin by walking along the vector 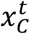.

We assumed that people could over, or underweight, a given leg when computing the sum – perhaps because they integrate evidence unevenly over time (Keung, Hagen, & Wilson, 2019). To model this suboptimality, we allowed 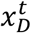 to be a *weighted* sum of the vectors from the first two legs:

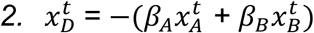

Where *β_A_* and *β_B_* denote the weights given to leg A and leg B respectively (Figure 1D). Combining the first two suboptimalities gives us Model 1, which includes noise and the possibility of over and underweighting the legs.

Of course, real participants are suboptimal and we modeled these suboptimalities in a number of different ways. First, people may not perfectly encode the vectors from the guided legs and/or may not perfectly implement the desired action, adding noise to the sum in equation 1. Thus, we assumed that the vector they actually walked 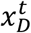 was sampled from a Gaussian distribution centered on 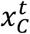, i.e.

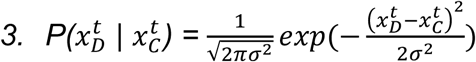

Where *σ*^2^ is the variance of the noise. Consistent with Weber’s law, we assumed this variance increased with the distance walked to match our finding of increased variance as a factor of distance walked in Experiment 2 (see Results, Figure 7B).

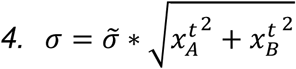

Finally, we allowed for the possibility that there may be sequential effects in our paradigm, i.e. there was an influence of previous trials on the current response. We modeled these sequential effects by including the vectors walked 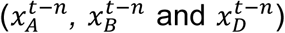 from past trials. For simplicity, we assumed that the effect of past trials decayed exponentially into the past (Lau & Glimcher, 2005), thus writing 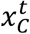 as

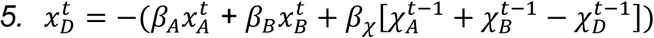

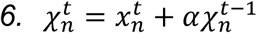

Where *χ*_n_ is a linear combination of the previous vectors, fitted with *α*, which ranges between 0 to 1, to capture the impact of prior trials. Thus, including the possible effect of past trials gave us Model 2.

#### Fitting the model

We fit the model using a maximum likelihood approach. In particular, we computed the log likelihood of the responses for each subject, as a function of model parameters:

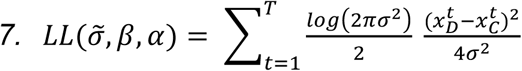

We then found the parameters that maximized the likelihood using Matlab’s fmincon function.

#### Simulating the models

To simulate the model, we used the parameter values fit for each subject to compute the mean 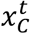 for each trial. To model the noise in each person’s choice, we perturbed the estimate of 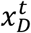 by isotropic Gaussian noise of mean 0 and variance *σ*^2^.

#### Encoding-Error Model

We recreated the Encoding-Error Model from Fujita et al. 1993. See Appendix A for more details. We used the same fitting and simulation method used for Model 1 and Model 2 with the exception of dividing the data for left and right-handed triangle to better accommodate the parameters of the Encoding-Error Model (see Klatzky et al. 1999).

#### Model Comparison Methods

We used two methods of model comparisons: 1) Penalized-Log-likelihood criteria’s Bayes Information Criterion (BIC) (Schwarz et al., 1978) and Akaike information criterion (AIC) (Akaike, 1974). Both express similar information about the generalizability of the model by penalizing for the number of free parameters. To test how meaningful our model comparisons results are in our task we also tested for model recovery. We did this by simulating each model with randomized parameter values and then fitting the models to the simulated data, allowing comparison of the AIC and BIC (see Wilson & Collins 2019 section 6 and Appendix B). We performed each simulation at the participant level and then subsequently compared BIC values by calculating exceedance probabilities, which measured how likely it is that the given model fits all of the data (Rigoux et al., 2014). This group level statistic is similar to AIC and BIC. Computed exceedance probabilities on our data as well as each model by simulating 100 times and comparing with the methods mentioned above. These methods are illustrated in Supplementary Figure 6 where the probability of the model fit for the simulated data ranges from 0 to1. The Exceedance Probability is calculated using SPM 12 spm_BMS function.

#### Bayes Factor Analyses

We included a Bayes Factor analysis for all statistical analyses (Rouder, Speckman, Sun, Morey, & Iverson, 2009). For results below our significance threshold (p<0.05), we used a Bayes Factor BF_***10***_ to indicate the degree of favorability toward the alternative hypothesis. For results that were not below our significance threshold, we employed the Bayes Null factor, BF_***01***_. Note that the larger the Bayes Factor, regardless of whether in favor of the alternative or null, the greater the evidence.

### Experiment 1

#### Participants

We tested a total of 26 participants (12m,14f), 4 (1m, 3f) of which were removed due to exceeding 25% of trials removed (see methods), Participants were tested on 7 different triangles described in detail in the methods (i.e., scalene, isosceles, right, equilateral, and isosceles-right). Estimates of sample size were based on the 12 participants used in Loomis et al. 1993 and in subsequent studies by Yamamoto 2013 et al. that employed a similar experimental design: as we were additionally testing a larger range of triangles, we thus approximately doubled the sample size.

#### Procedure

We outline the basic set up for triangle geometry in Figure 1E, which shows the stacked triangle templets, with a constant 10m leg C’ (unguided leg), while manipulating the angle. The 7 triangle configurations are shown in Supplementary Table 1A, with 3 scalene, 1 isosceles, 1 right, 1 equilateral, and 1 isosceles-right. To keep leg C’ at 10m across all 7 triangles, we employed different leg A’ and leg B’ sizes to accommodate the different angles. There were 28 trials, in which 14 of them were left-handed (subjects only made left turn) and 14 right handed (subject only made right turns). We did this to avoid any advantages for right vs. left turns during the task.

In Experiment 1, as part of ensuring the compliance and efficacy of the hand-held controllers in following the guided legs, we compared with a condition in which participants walked the guided legs on half the trials using a visual beacon. In this situation, participants saw a large red monolith that they walked to while receiving feedback from the handheld controllers. It is important to emphasize that the vision-guided trials were only present for the *guided* legs and were simply to ensure that participants accurately encoded the guided legs before performing the unguided legs.

### Experiment 2

#### Participants

We tested a total of 21 participants (9m,11f), 3 (1m 2f) of which did not complete the experiment, with additional 1 female participant removed from the analysis for exceeding 25% trials below criterial performance. Given the longer distances in Experiment 2, participants were allowed to take a break, but only at the end of a trial. About 50% of participants took a break at some point during the experiment.

#### Procedure

Here, we employed scalene triangles with different length perimeters to allow us to manipulate distance while keeping angle relatively constant, testing 5 different triangle sizes. Figure 1F shows the stacked triangle templates we employed with constant internal angles but varying in size. The triangle configurations are shown in Supplementary Table 1B, with 15m, 25m, 127m, 253m, and 506m perimeters. There were 30 trials, with 15 of them left-handed (participants only made left turn) and 15 right-handed (participants only made right turns). Unlike Experiment 1, there were no vision trials. Due to testing longer distances and wanting to avoid fatigue, we limited the number of trials for the longest distance triangles. The distributions of trials were 10 for the15m triangle, 10 for the 25m triangle, 8 for the 127m triangle, 4 for the 253m triangle, and 2 for the 506m triangle.

All data files are available at: github.com/sharootonian/PA-TCT

## Acknowledgements.

The Authors are gratful to E. Erlenbach for helping during data collection. Research supported by grants from NSF Division of Behavioral and Cognitive Sciences [BCS-1630296] awarded to Arne Ekstrom.

## Supplementary figures

**Supplementary Figure 1.**
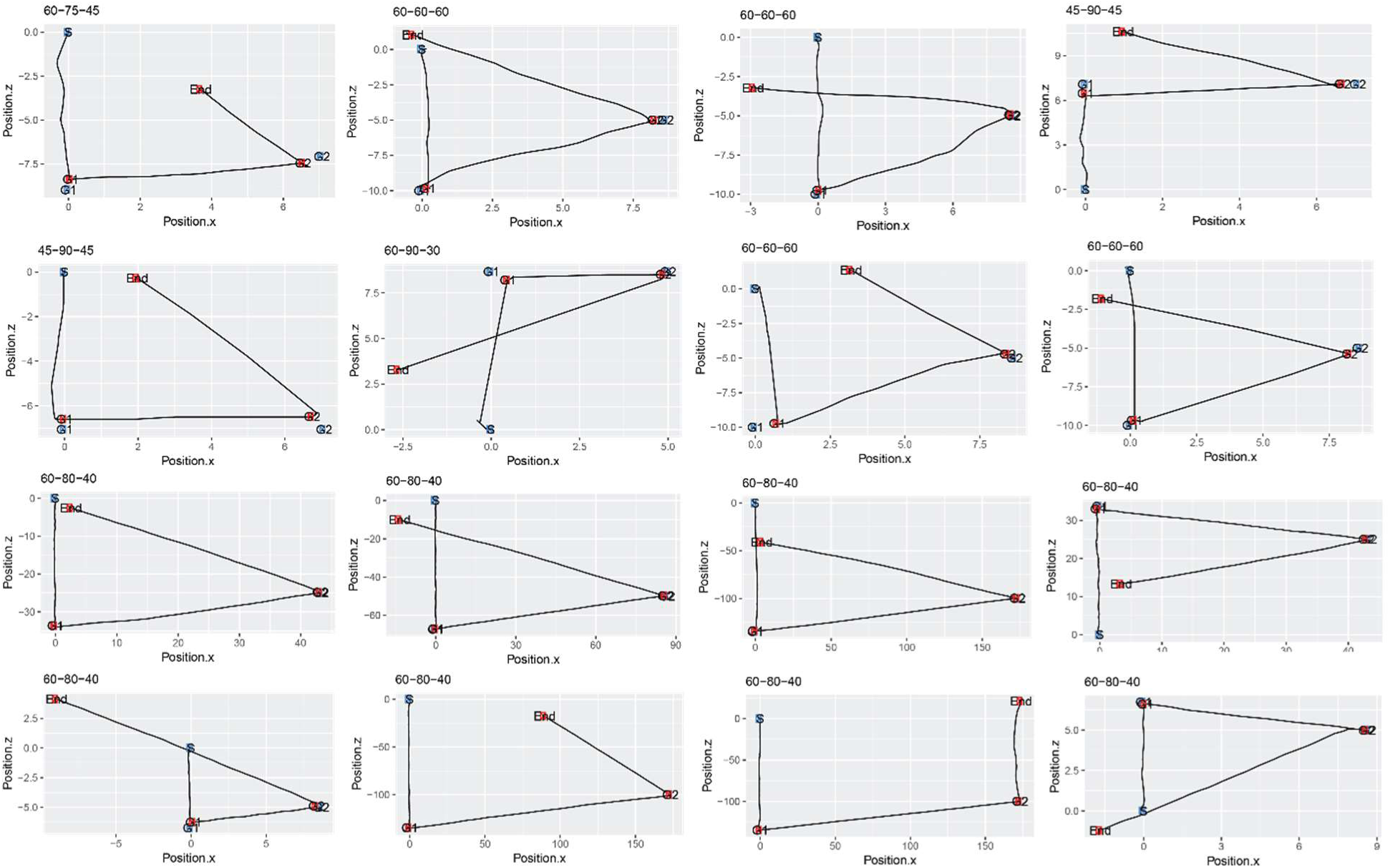
: Raw trials from experiment 1 (top 8) and experiment 2 (bottom 8).

**Supplementary Figure 2.**
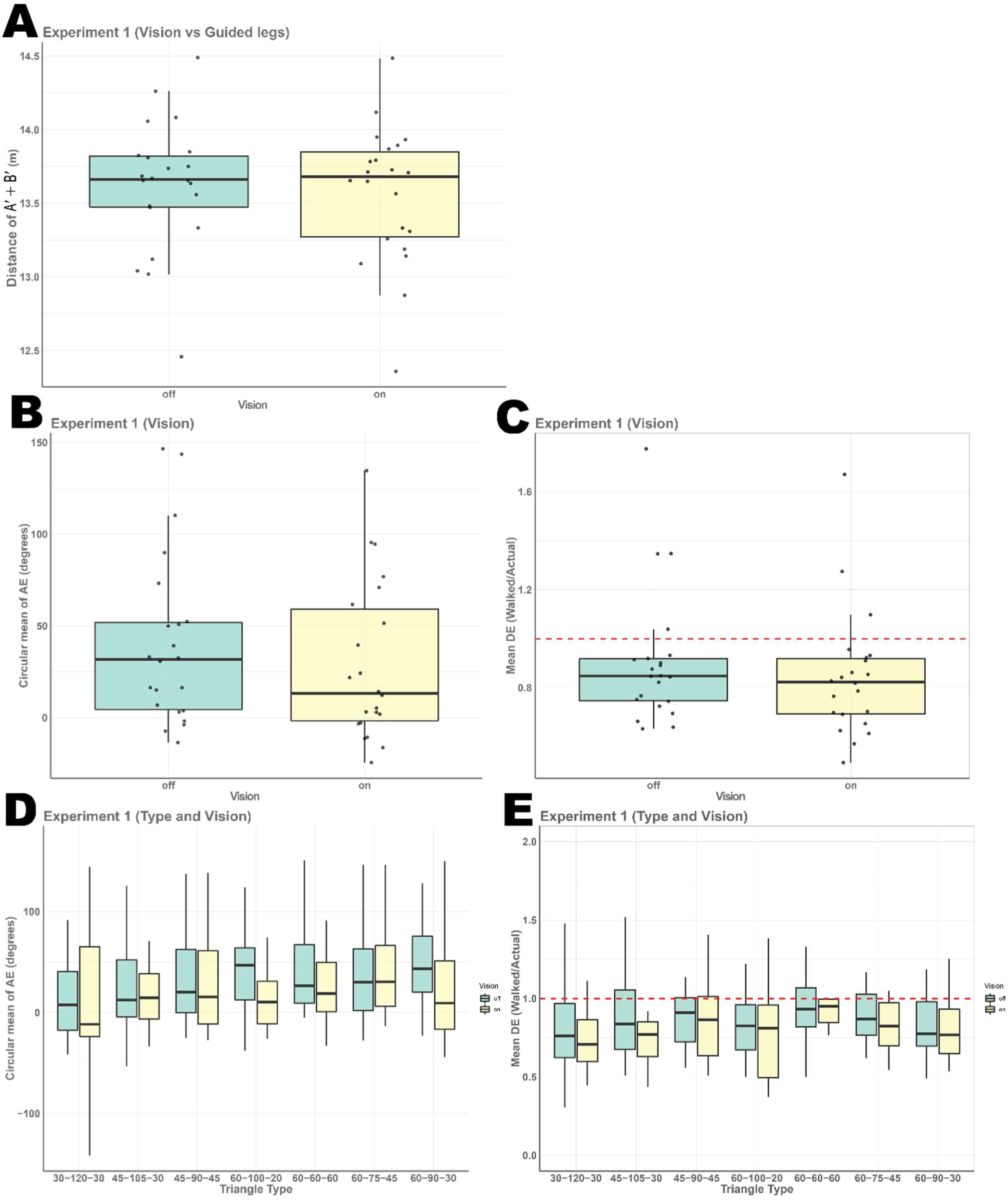
: (A) Combined distance walked during guided legs during vision on and vision off trial, showing now differences (t(21)=1.09, p=0.288, Cohen’s d=0.336 and BF_01_>3) (B) Angle error from experiment 1, showing a small but significant difference between vision on and off condition (t(21)=2.46, p<0.022, Cohen’s d=0.248 and BF_10_=2.54) (C) Distance error from experiment 1, showing a significant difference between vision on and off condition (t(21)=2.71, p<0.013, Cohen’s d=0.232 and BF_10_=3.94). (D) Angle error from experiment 1, ANOVA significant for triangle type F(6,21)=2.9, p<0.01, η^2^ =0.058 BF_10_=1.72. and Vision F(1, 21)=4.9, p<0.026, η^2^=0.016 BF_10_=1.16, but not for the interaction between Type and Vision F(6, 21)=1.454, p=0.194, η^2^ =0.029 BF_10_=0.432. (E) Distance error from experiment 1, ANOVA significant for triangle type F(6, 21)=5.7, p<0.1.33e-5, η^2^=0.109 BF_10_>10 and r Vision F(1, 21)=8.2, p<0.004, η^2^ =0.026 BF_10_>4, but not for the interaction between Type and Vision F(6, 21)=0.199, p=0.976, η^2^=0.004 BF_10_>10.

**Supplementary Figure 3:**
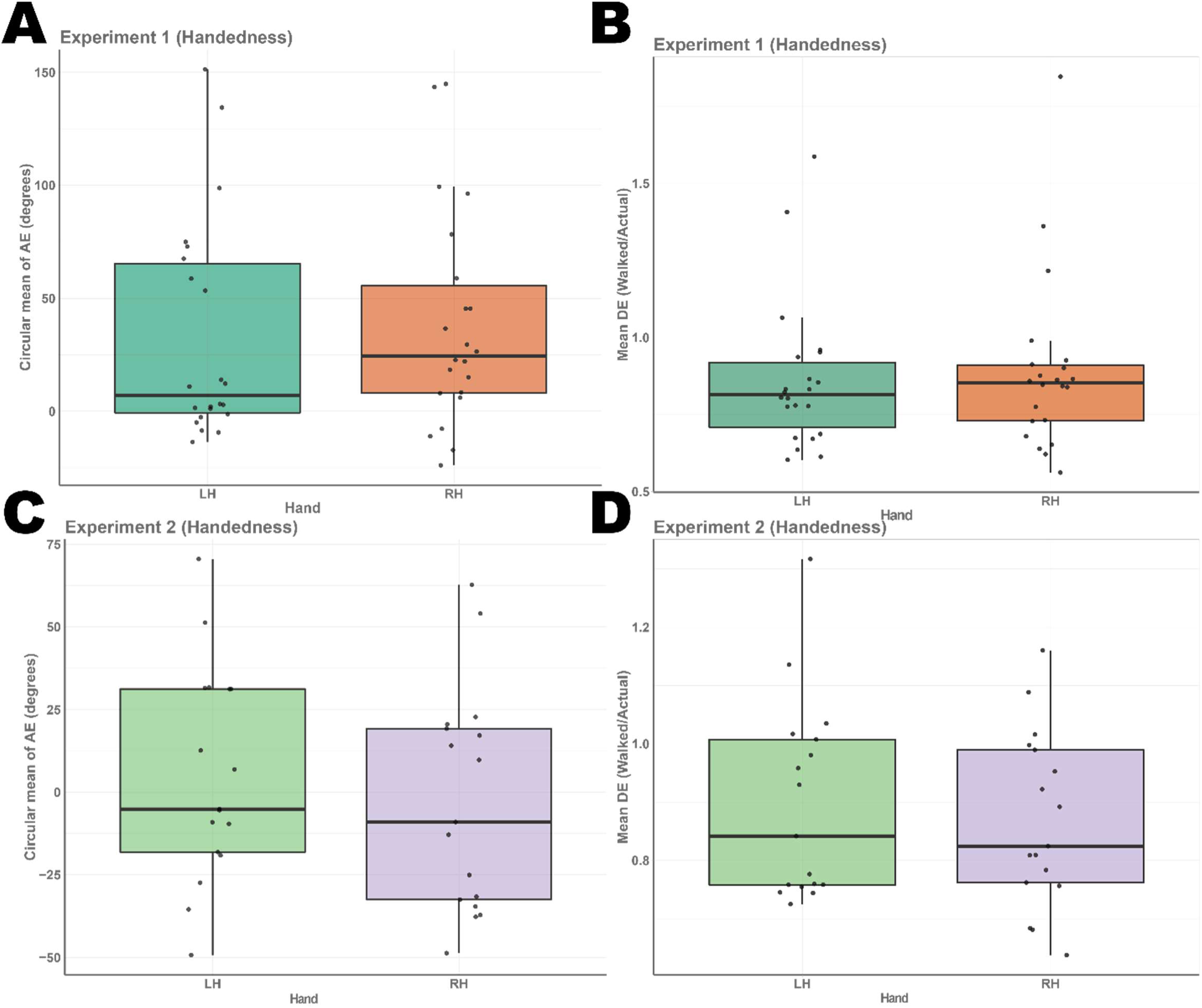
(A) Angle error from experiment 1 which showed no difference between left and right-handed triangles (t(21)=0.7, p=0.485, Cohen’s d=0.118 and BF_01_>3). (B) Distance error from experiment 1, which showed no difference between left and right-handed triangle (t(21)=1.136, p=0.268, Cohen’s d=0.103 and BF_01_=2.53). (C) Angle error from experiment 2, again showing no difference between left and right-handed triangle (t(16)=1.51, p=0.151, Cohen’s d=0.245 and BF_01_=1.55). (D) Distance error from experiment 2, which showed no difference between left and right-handed triangle (t(16)=0.724, p=0.4797, Cohen’s d=0.176 and BF_01_=3.188).

**Supplementary Figure 4:**
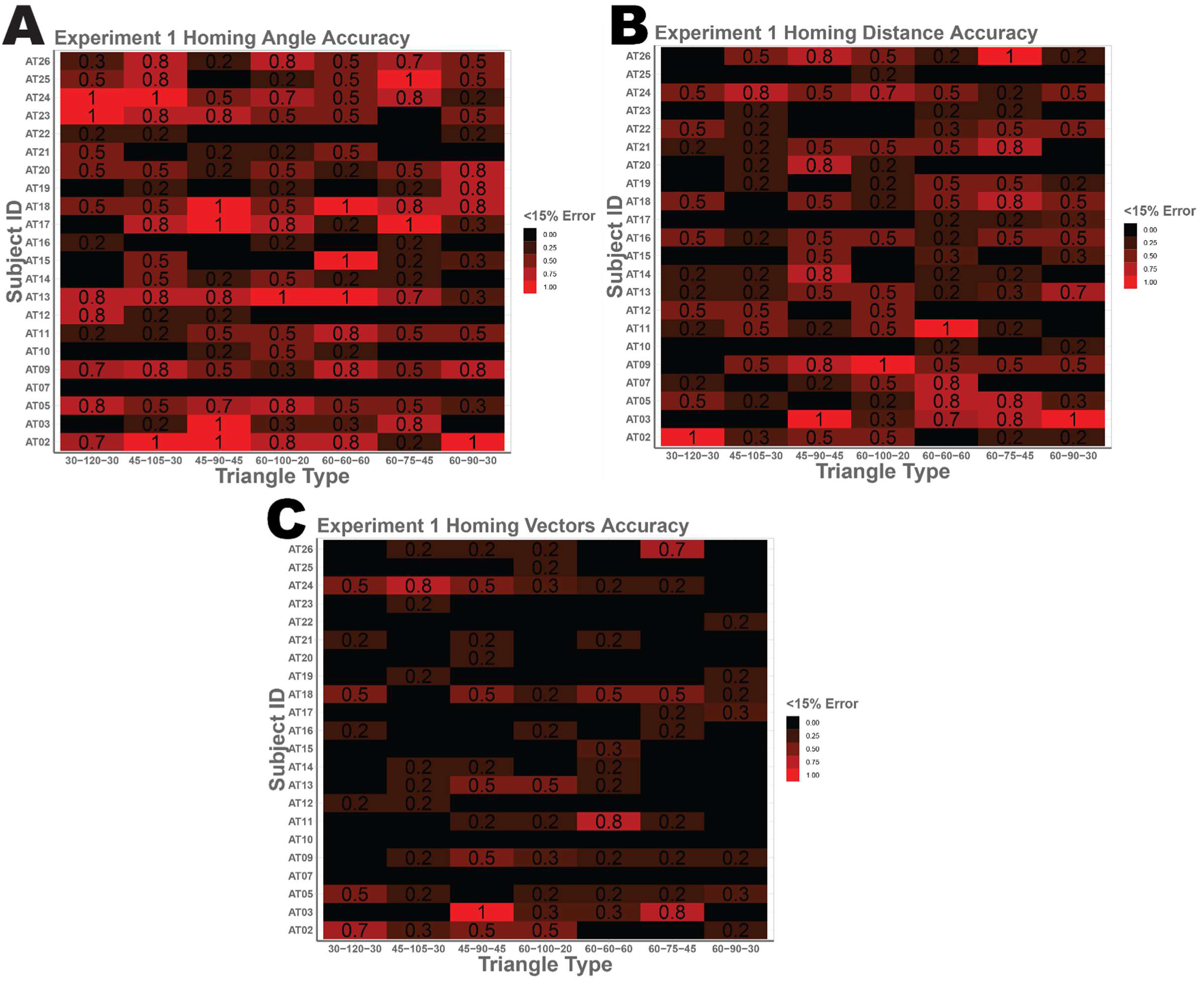
Raster plot (A) showing the percentage of responses with less than 15% angle error (ranging from −27° to 27°) for triangle type (x-axis) and participants (y-axis). Participants are 281.39% more likely to have <15% angle error in their unguided leg then <15% total error (angle and distance). (B) percentage of responses with less than 15% distance error (8.5m to 11.5m). Participants are 208.14% more likely to have <15% distance error in there unguided leg then <15% total error (angle and distance). (C) percentage of responses with less than 15% angle error (ranging from −27° to 27°) and 10% distance error (8.5m to 11.5m). In (C) we can see that all of participant AT03’s responses for triangle 45-90-45 are less then 15% error for both angle and distance error. And 80% for equilateral triangle (60-60-60) for participant AT11.

**Supplementary Figure 5:**
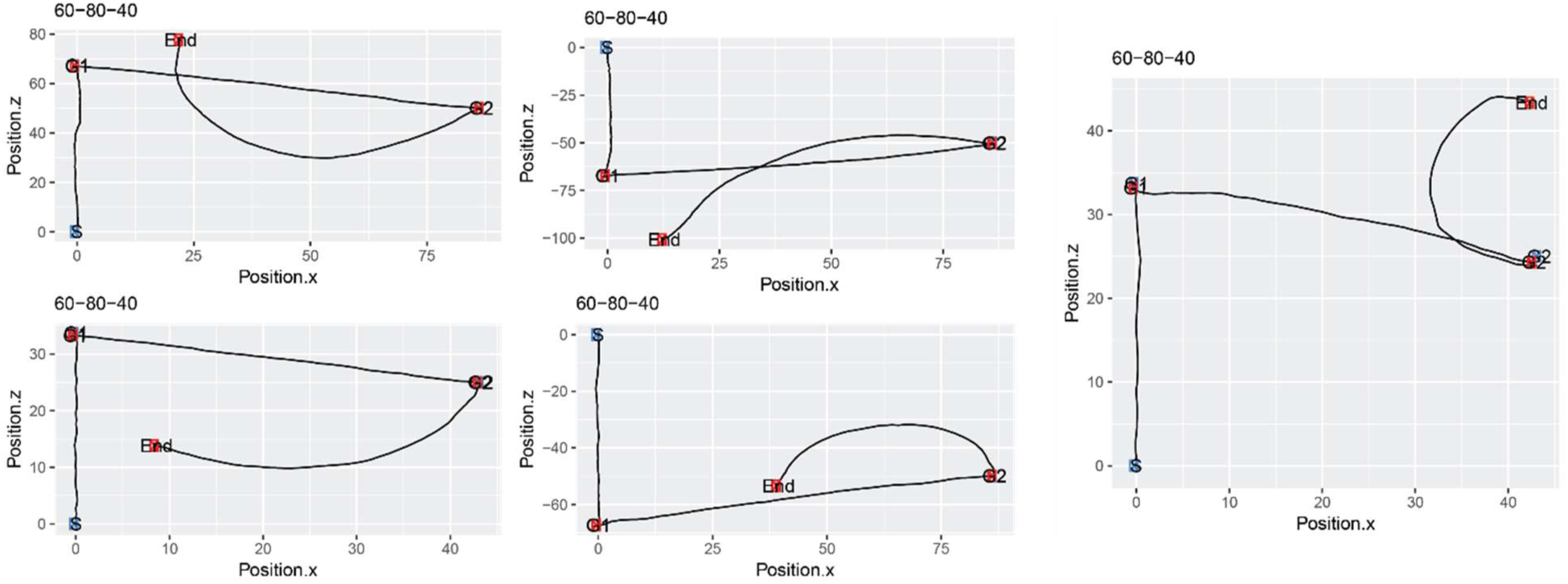
Raw trials that were removed due to circular pathing.

**Supplementary Figure 6:**
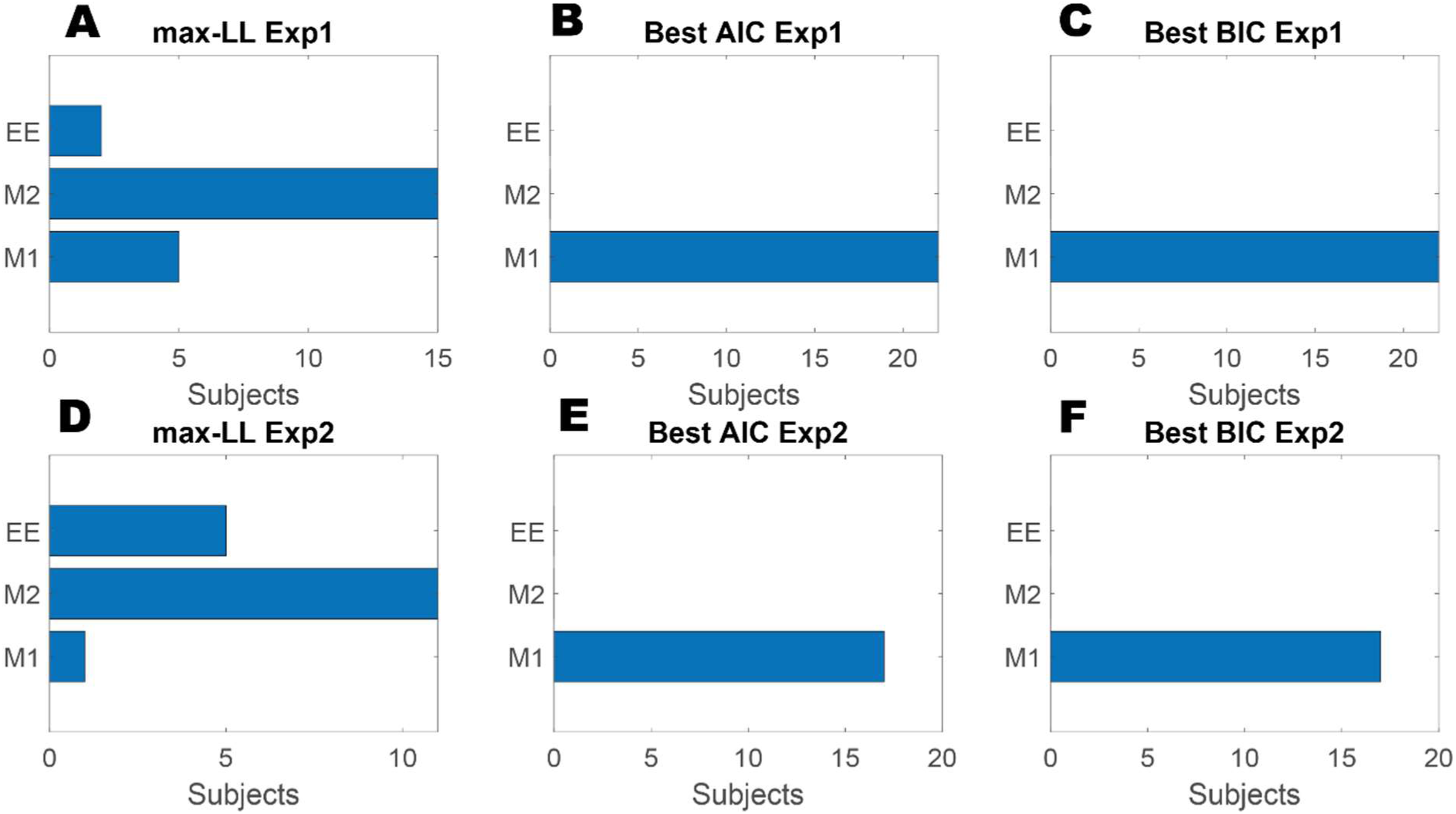
Comparing model fitting of the individual participant’s data. A) Shows best model fit (highest loglikelihood values) for each subject in experiment 1. B&C) Lowest AIC and BIC values across the 3 models for each subject in experiment 2. D) Shows best model fit (highest loglikelihood values) for each subject in experiment 2. E&F) Lowest AIC and BIC values across the 3 models for each subject in experiment 2

**Supplementary Figure 7:**
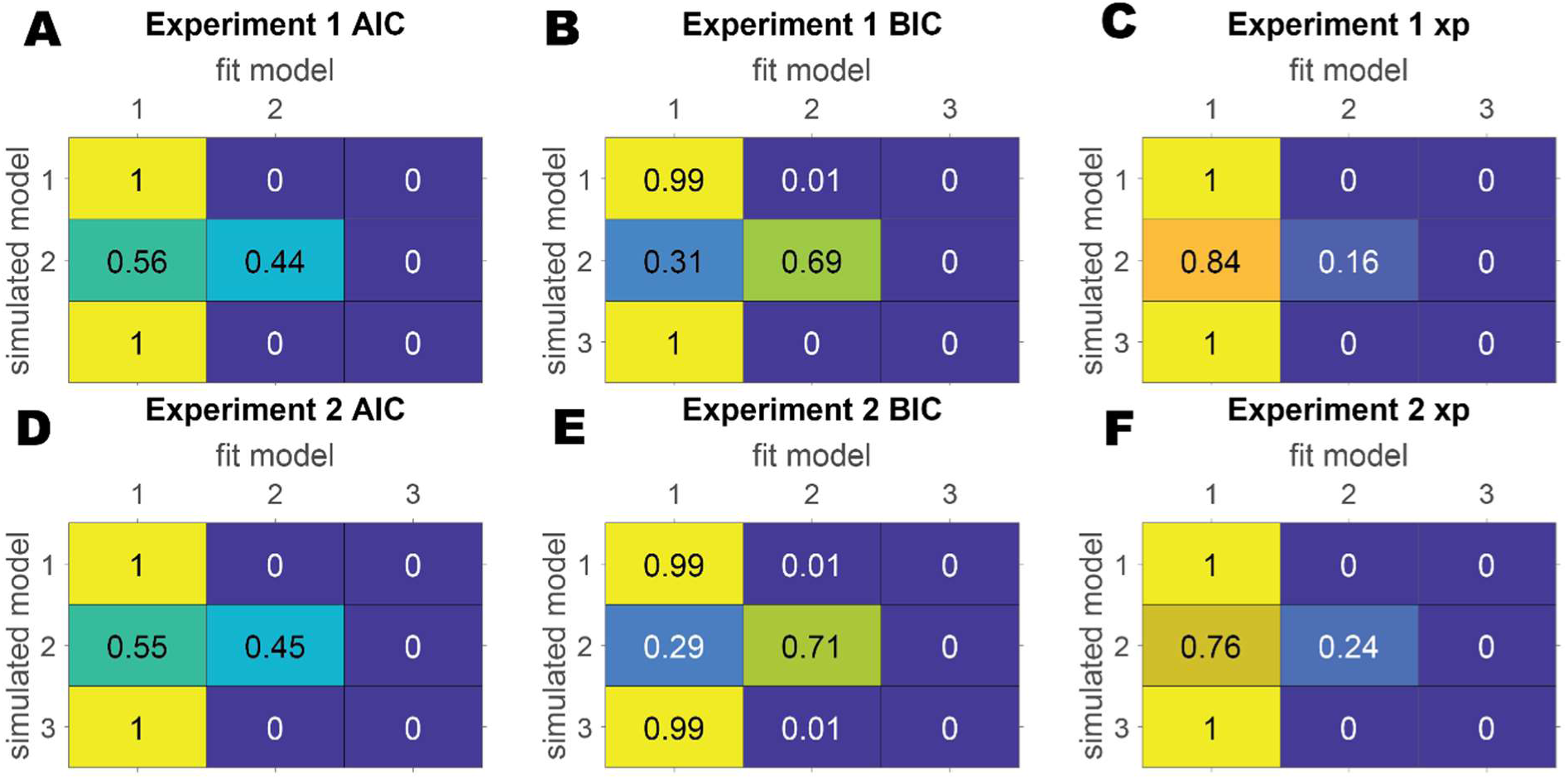
Model recovery confusion matrices where 1 in column and row represents Model 1, 2 represents Model 2 and 3 represents Encoding-Error Model. Probability ranges from 0 to 1. (A & B) Show best AIC and BIC for Experiment 1 respectively. The higher value in the diagonal shows better model recovery from this experiment. We see that the Encoding-Error does not fit its own simulated data well. (C)The Exceedance Probability for Experiment 1. (D & E) Show best AIC and BIC for Experiment 2 respectively. Again, we see Encoding-Error does not fit its simulated data well. (C)The Exceedance Probability for Experiment 2.

**Supplementary Figure 8:**
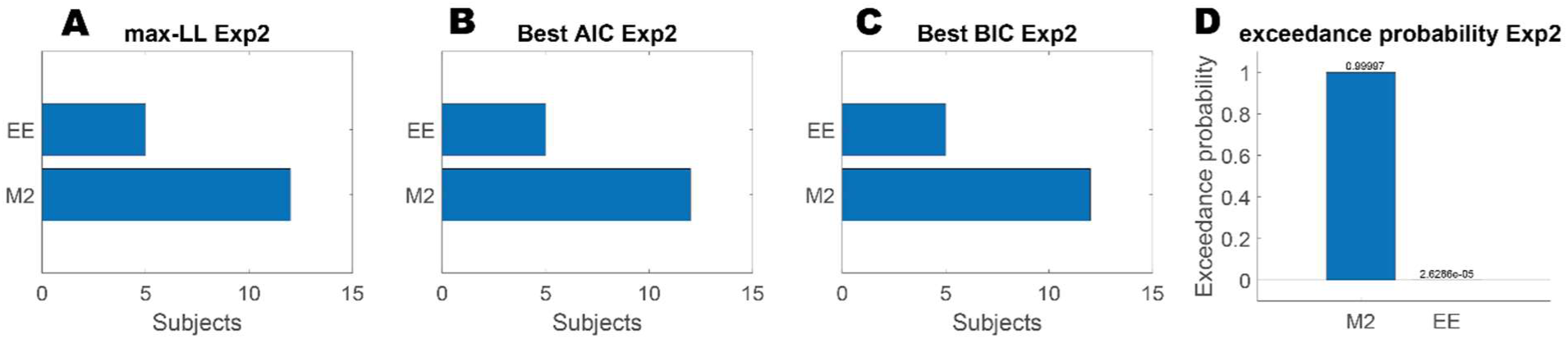
Comparing model fitting of the individual participant’s data. A) Shows best model fit (highest loglikelihood values) for each subject in experiment 2. B&C) Lowest AIC and BIC values across the 3 models for each subject in experiment 2. D) Shows the exceedance probability of each model for experiment 2.

**Supplementary Figure 9:**
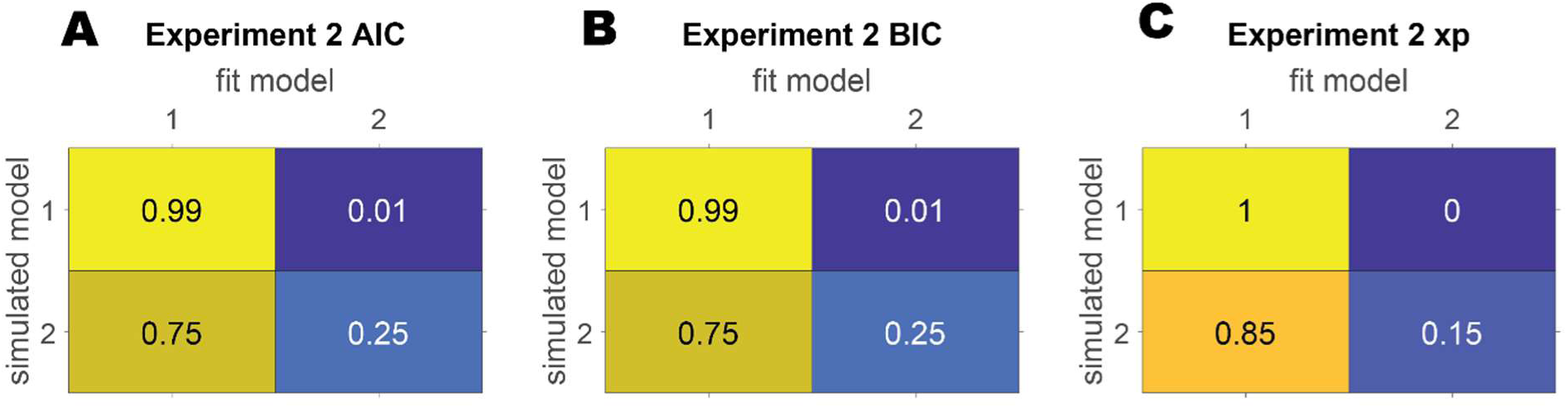
Model recovery confusion matrices where 1 in column and row represents Model 2 and 2 represents Encoding-Error Model. Probability ranges from 0 to 1. (A & B) Show best AIC and BIC for Experiment 2 respectively. We see Encoding-Error does not fit its own simulated data well. (C)The Exceedance Probability for Experiment 2.

## Supplementary Table 1

**Supplementary Table 1A:** Showing the configuration of each triangle using in experiment 1.

Experiment 1
| <BC | <AB | <CA | Side A (m) | Side B (m) | Side C (m) | Perimeter (m) | Type |
| --- | --- | --- | --- | --- | --- | --- | --- |
| 60 | 100 | 20 | 8.793852 | 3.472964 | 10 | 22.267 | scalene |
| 30 | 120 | 30 | 5.773503 | 5.773503 | 10 | 21.547 | isosceles |
| 45 | 105 | 30 | 7.320508 | 5.176381 | 10 | 22.497 | scalene |
| 60 | 90 | 30 | 8.660254 | 5 | 10 | 23.660 | right |
| 60 | 75 | 45 | 8.965755 | 7.320508 | 10 | 26.286 | scalene |
| 45 | 90 | 45 | 7.071068 | 7.071068 | 10 | 24.142 | isosceles-right |
| 60 | 60 | 60 | 10 | 10 | 10 | 30.000 | equilateral |

**Supplementary Table 1B:** Showing the configuration of each triangle using in experiment 2.

| <BC | <AB | <CA | Side A (m) | Side B (m) | Side C (m) | Perimeter (m) | Type |
| --- | --- | --- | --- | --- | --- | --- | --- |
| 40 | 80 | 60 | 4.042 | 5.299 | 6 | 15.193 | scalene |
| 40 | 80 | 60 | 6.736 | 8.832 | 10 | 25.321 | scalene |
| 40 | 80 | 60 | 33.682 | 44.163 | 50 | 126.604 | scalene |
| 40 | 80 | 60 | 67.365 | 88.327 | 100 | 253.209 | scalene |
| 40 | 80 | 60 | 134.73 | 176.653 | 200 | 506.418 | scalene |

## Footnotes

1 Klatzky et al. 199 state: “The assumption of immutable encoding seems, a priori, to be doubtful. Encoding of pathways on the scale of tens of meters is unlikely to use the same mapping as is used for pathways on the scale of under 10 m (p. 35)

